# Live-cell imaging of circadian clock protein dynamics in CRISPR-generated knock-in cells

**DOI:** 10.1101/2020.02.28.967752

**Authors:** Christian H. Gabriel, Marta del Olmo, Amin Zehtabian, Silke Reischl, Hannah van Dijk, Barbara Koller, Astrid Grudziecki, Bert Maier, Helge Ewers, Hanspeter Herzel, Adrian E. Granada, Achim Kramer

## Abstract

The current model of the mammalian circadian oscillator is predominantly based on data from genetics and biochemistry experiments, while the cell biology of circadian clocks is still in its infancy. Here, we describe a new strategy for the efficient generation of knock-in reporter cell lines using CRISPR technology that is particularly useful for lowly or transiently expressed genes, such as those coding for circadian clock proteins. We generated single and double knock-in cells with endogenously expressed PER2 and CRY1 fused to fluorescent proteins, which allowed to simultaneously monitor the dynamics of CRY1 and PER2 proteins in live single cells. Both proteins are highly rhythmic in the nucleus of human cells with PER2 showing a much higher amplitude than CRY1. Surprisingly, CRY1 protein is nuclear at all circadian times indicating the absence of circadian gating of nuclear import. Furthermore, in the nucleus of individual cells CRY1 abundance rhythms are phase-delayed (∼5 hours), and CRY1 levels are much higher (>6 times) compared to PER2 questioning the current model of the circadian oscillator. Our knock-in strategy will allow the generation of additional single, double or triple knock-in cells for circadian clock proteins, which should greatly advance our understanding about the cell biology of circadian clocks.

## INTRODUCTION

To stay in synchrony with environmental cycles, most living organisms developed endogenous clocks, which regulate the circadian (∼24 h) rhythms of molecular, physiological, and behavioral functions. The molecular basis of these circadian clocks is a gene-regulatory network with transcription-translation feedback loops driving cell-autonomous oscillations in most mammalian tissues. According to the current model, the heterodimeric transcription factor CLOCK:BMAL1 mediates the expression of rhythmically transcribed genes by binding to E-box enhancer elements in their promoters (*1*). Among those genes are the canonical repressors PER1-3 and CRY1-2, which inhibit CLOCK:BMAL1 transcriptional activity after a delay of several hours and, thereby, their own expression (*2, 3*). After the regulated degradation of PERs and CRYs, the transcription factor complex is released from repression, and a new cycle can start (*4–6*). Similar to BMAL1-KO animals, PER- or CRY-deficient mice are behaviorally arrhythmic, emphasizing the importance of each protein family for the integrity of the molecular clock (*7, 8*).

PER and CRY proteins physically interact (*9–11*), and there is evidence that this interaction affects their subcellular localization. Genetic studies show that CRYs do not accumulate in the nucleus of *Per1/2* double knock-out cells and, similarly, PERs are almost exclusively cytoplasmic in cells lacking both CRY proteins (*12*) suggesting that the presence of each family is necessary for proper PER and CRY protein localization. This is supported by overexpression studies, in which CRY1 is accelerating PER2 nuclear import dynamics in human cells (*13*). In *Drosophila*, analogs of PERs and CRY, dPER and TIM, first accumulate in the cytoplasm when overexpressed and after a delay of several hours translocate into the nucleus together (*14*). Although similar models were proposed for the mammalian system (*15*), no circadian differences in subcellular localization of PER2 were observed in cells from PER2-Venus knock-in mice (*16*), thus questioning this analogy.

Recent biochemical studies with mouse liver lysates suggest that during the repressive phase, essentially all nuclear PER and CRY proteins are coordinated together in one large repressive complex, with only a minor amount of CRY1 monomers detectable (*17*). Again, double knock-out of either *Per1/2* or *Cry1/2* completely prevented the formation of this repressive complex.

Most of the current knowledge of PER and CRY protein dynamics resulted either from biochemical data with mixed lysates of many thousand cells, or from single-cell imaging of over-expressed fluorescent tagged fusion proteins (*12, 13, 17, 18*). Both approaches have clear limitations: Population sampling - e.g. cell fractionation followed by Western Blot, chromatography or immunoprecipitation - not only conceals spatial information but also suffers from much reduced temporal resolution. Most importantly, however, population sampling averages signals from thousands of cells thereby masking individual cell properties (e.g. regarding circadian period, phase and amplitude) and degree of noise. While fluorescent tagged proteins constitute an outstanding tool to monitor protein expression and localization in individual living cells, overexpression of PER-and CRY-proteins in most cases disrupts the circadian oscillator and data from such experiments have to be conceived with caution (*19, 20*).

Such limitations can be overcome by incorporating a fluorescent tag directly into the proteins’ genomic locus. In this case, expression dynamics and level of the resulting fusion protein often remain similar to the wild-type protein and the clock stays intact. Indeed, the PER2-Luciferase and the PER2-Venus knock-in mice – in which PER2 is tagged at the genomic level with a luciferase or a yellow fluorescent protein, respectively – enabled analysis of PER2 protein oscillations on a single cell level without compromising the oscillator (*16, 21*). In contrast, expression and localization dynamics of endogenously expressed CRY proteins in live cells have not been reported yet, due to the lack of similar knock-in models. Furthermore, differences between the murine and primate circadian oscillator create a need for human cellular models (*22*). Thus, this lack of model motivated us to create human cell lines that express fluorescence tagged versions of PER and CRY proteins from the respective endogenous loci.

While targeted introduction of DNA into the genome of a somatic cell used to be extremely inefficient and – if possible at all – laborious, the discovery and development of CRISPR/Cas9 based genome editing changed the game (*23, 24*). In short, targeted Cas9-mediated DNA double strand breaks are – among other possible outcomes – eliminated by the endogenous homology directed repair (HDR) pathway, which can be hijacked to introduce an exogenous donor sequence (such as a fluorescent protein tag) into the locus (reviewed in (*25*)). Although Cas9-induced double strand breaks greatly stimulate the integration of such a homologous donor, the rate of targeted integration is usually still low and depends on many parameters, such as cell type, transfection efficiency and length of the integrated sequence. In addition, existing strategies to enrich for the desired cells are prone to fail when targeting lowly and/or transiently expressed genes.

Here, we report a new strategy to efficiently knock-in fluorescent reporter proteins into the genomic locus of the lowly expressed circadian proteins PER2 and CRY1 and applied it to human cells. We generated single and double knock-in cells with intact circadian clocks, which allowed us to monitor the dynamics of CRY1 and PER2 fusion proteins in live single cells. We found that CRY1 protein is mainly nuclear at all circadian times suggesting absence of circadian gating of nuclear import. Furthermore, CRY1 accumulates phase-delayed and to much higher (>6 times) levels compared to PER2 protein in the nucleus of individual cells questioning the current model of the circadian oscillator.

## RESULTS

### Strategy for CRISPR/Cas9-mediated knock-in

To insert reporter protein tags into the *PER2* and *CRY1* genomic loci of human cells, we conceived a knock-in strategy for lowly expressed genes. Thereby various tags including mClover3 (*26*) and mScarlet-I (*27*), bright monomeric green or red fluorescence proteins (FP), respectively, as well as firefly luciferase were aimed to be integrated into these loci. In the following, we exemplarily outline the vector design as well as the screening strategy for generating knock-in cells that express PER2 C-terminally tagged to mScarlet-I from the endogenous *PER2* locus.

To this end, we intended to integrate the FP-coding sequence directly upstream of the STOP codon of *PER2* (**Fig. 1A**). Hence, we designed sgRNAs that target the Cas9 to introduce double strand breaks very close (<60 bp) to the STOP codon sequence. The donor vector that is to be integrated during HDR, contained the coding sequence of mScarlet-I flanked by *PER2* sequences (∼800 bp) homologous to those directly upstream and downstream of the STOP codon. A C-terminal 6xHis/FLAG-tag (HF-tag) and a new STOP codon for the fusion protein followed downstream of mScarlet-I (**Fig. 1A**). Importantly, we mutated the PAM sequences for the sgRNAs in the donor vector to prevent Cas9 from cutting donor vector and the edited gene.

**Figure 1:**
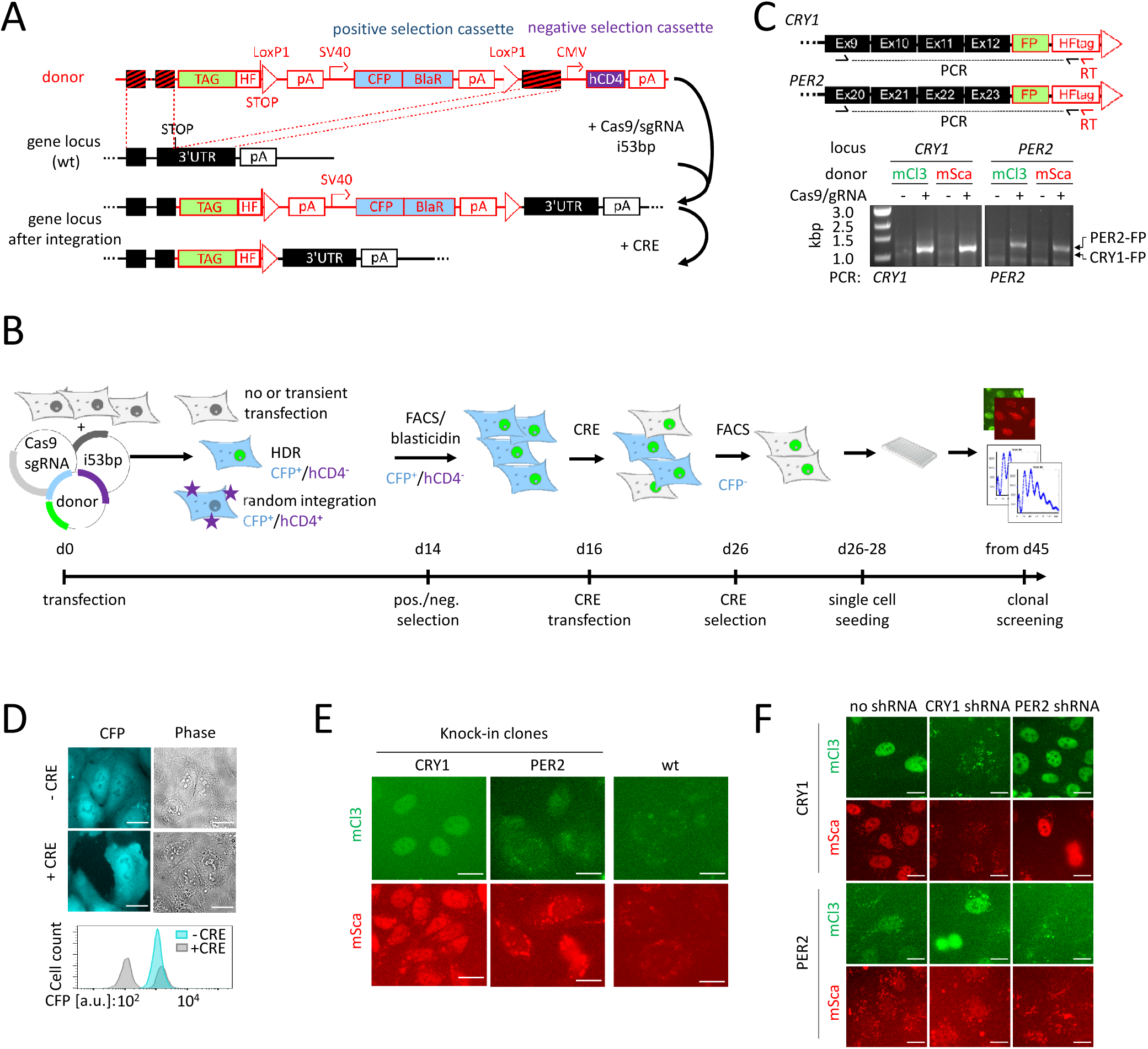
CRISPR/Cas9-mediated generation of clock protein knock-in reporter cells. **(A)** Donor plasmid design and genome editing strategy: The tag (e.g. fluorescent protein) to be integrated and a floxed positive selection cassette (CFP + blasticidin-resistance) are flanked by arms which are homologous to the genomic target region. When Cas9/sgRNA-mediated DNA double strand breaks are repaired by HDR, tag and positive selection marker are integrated into the target region. The negative selection cassette (hCD4, a cell surface protein exclusively expressed on immune cells) is only integrated into the genome by unwanted random integration of the whole donor plasmid. **(B)** Selection strategy. Cells are transfected with Cas9, sgRNA, i53bp (see text) and donor plasmid. Stable transfectants are selected by blasticidin selection and fluorescence activated cell sorting (FACS) of CFP positive cells, while unwanted hCD4 positive cells are depleted. Subsequently, cells are transiently transfected with CRE recombinase to remove the positive selection cassette from the genomic locus, and only CFP negative cells are clonally expanded and screened for successful knock-in. **(C)** Chimeric mRNA was detected in selected batch cultures by RT-PCR using a RT- and a reverse primer specific to the insertion and a gene specific forward primer. **(D)** Loss of CFP expression after removal of the positive selection cassette monitored by microscopy and flow cytometry. **(E)** Fluorescence microscopy images of successful knock-in clones. **(F)** Indicated knock-in cells were either left untreated or transduced with shRNA targeting either *CRY1* or *PER2*. Images were acquired 10 h after synchronization. Scale bars: 20 µm. mCl3 = mClover3, mSca = mScarlet-I, FP = fluorescent protein, HF: His-tag/FLAG-tag

Due to the general low efficiency of HDR, it was expected that only a small fraction of cells transfected with Cas9, gRNA and donor vector would integrate the donor sequence into the genome, while the majority would be transiently transfected. Several strategies have been described to enrich for cells with successful HDR. For example, the donor can be designed in such a way that integration results in the co-expression of a marker protein, such as a fluorescent protein or an antibiotic resistance. Positive selection by fluorescence-activated cell sorting (FACS) or antibiotic treatment then allows the depletion of all cells without donor integration and most cells with random, non-targeted integration that often does not lead to marker expression.

A drawback of such methods is that they have limits for editing of lowly and/or transiently (e.g. rhythmically) expressed genes (such as *PER2*), because marker expression strength correlates with (in this case low) expression of the target gene. Thus, correctly edited cells often cannot be identified using FACS, because fluorescence levels may not exceed cellular auto-fluorescence. Similarly, low or transient expression of an antibiotic resistance marker may not be sufficient to confer the cells resistance to the respective antibiotic. In both cases, correctly edited cells may be co-depleted during ‘enrichment’ steps.

To circumvent these limitations for the low copy number and transiently expressed *PER2* (estimated protein molecules between 0 and 10.000 per cell (*28*)), we placed the selection marker in a separate expression cassette downstream of the FP-STOP-codon. Robust marker expression is driven by an SV40 promoter thus independently of the expression of *PER2*. As a marker, we used a self-cleaving CFP-BlaR fusion protein that allows for selection either by blasticidin treatment or by FACS of cyan fluorescent protein (CFP) positive cells. To ensure expression of the PER2 fusion protein, we introduced a poly(A) transcriptional termination site between the STOP codon and the promoter of the positive selection cassette.

This strategy enables enrichment of cells with genomic integration of the donor by elimination of non- and transiently transfected cells. However, it cannot discriminate whether cells have integrated the positive selection cassette via HDR at the correct genomic location or whether random insertion of the vector occurred anywhere in the genome, as the SV40 promoter will always drive marker expression. To address this issue, we reasoned that HDR will only integrate the sequence between the homologous arms, while random insertion would more likely integrate the whole vector into the genome. To exploit this mechanistic difference for further enrichment of correctly edited cells, we placed a negative selection cassette into the donor vector outside the homologous arms. This cassette expresses hCD4-extracellular domain and will only be integrated into the genome if the vector is randomly integrated, but not upon HDR. Hence, cells with random integration events can be efficiently eliminated by FACS or MACS.

After HDR-mediated integration of the FP and the positive selection cassette, transcription of the *PER2*-*FP*-mRNA would terminate at the incorporated poly(A)-site, resulting in a very short 3’-UTR. Since the endogenous 3’-UTR downstream of the positive selection cassette may include posttranscriptional regulation sites, we flanked the poly(A)-site and the positive selection cassette by LoxP sites. Thereby, transfection with a CRE expression plasmid will result in removal of the positive selection cassette and restoration of the endogenous 3’-UTR, which can be followed by loss of CFP expression.

To summarize, we designed a donor vector with essentially four features: (i) Homology arms plus FP for HDR-mediated editing of target gene. (ii) A positive selection cassette with an independent promoter and poly(A) site driving marker gene expression to select for cells with genomic donor vector integration. Thereby, low or transient expression does not interfere with the selection process. (iii) A negative selection cassette placed outside the homology regions to allow for depletion of cells with random integration of the donor vector. (iv) LoxP sites for removal of the positive selection cassette by transient CRE activity, restoring the genomic locus to essentially an endogenous constitution.

### Generation of knock-in cells

To create knock-in U-2 OS cells expressing PER2 or CRY1 C-terminally fused to the red-fluorescent protein mScarlet-I or to the green fluorescent protein mClover3 from their endogenous promoters, we transfected U-2 OS wild-type cells with three different plasmids: an spCas9/sgRNA expression plasmid, the donor plasmid, and a plasmid expressing an inhibitor of the 53-binding protein, which enhances HDR efficiency by suppressing NHEJ (*29*) (**Fig. 1B**). Depending on pre-determined sgRNA efficiency (not shown), either a single sgRNA (*CRY1*) or a mixture of different sgRNAs targeting the same region (*PER2*) were used.

Two weeks after transfection, the desired cell population was enriched by FACS for CFP^+^/hCD4^−^ cells, thereby depleting non- or only transiently transfected cells as well as cells that had undergone random integration of the donor plasmid. To check for successful integration of the fluorophores, we analyzed the selected cell populations for the presence of chimeric mRNA (target gene and fluorescence tag) by RT-PCR (**Fig. 1C**). As expected for specific knock-in, bands corresponding to the desired chimeric mRNA were detected in cell populations transfected with donor plasmid plus Cas9/sgRNA, but not in control populations transfected with donor plasmid only, indicating that a substantial number of cells of the population had undergone the intended integration events.

To eliminate the floxed positive selection cassette from the edited gene loci, we transfected the selected cell population with a CRE recombinase expression plasmid (**Fig. 1A**). Successful elimination was monitored by the loss of CFP fluorescence (**Fig. 1D**). Typically, 20-70 % of the cells became CFP-negative within 10 days after transfection without any selection step, and the remaining CFP-positive cells were removed by FACS.

Single cells were seeded into 96 well plates and after 2-3 weeks, clonal colonies were inspected using fluorescence microscopy. In total, 13 out of 31 examined potential *CRY1* knock-in clones showed a nuclear fluorescence consistent with data from CRY1 overexpression (*13*) (**Fig. 1E, Supplementary Fig. S1A**). For *PER2*, 7 out of 33 examined clones exhibited more diffuse fluorescence patterns exceeding auto-fluorescence (**Fig. 1E, Supplementary Fig. S1A**). Fluorescence signals of the other clones were not distinguishable from wild-type cells that served as negative controls (compare **Fig. 1E**, right panel). Notably, overall fluorescence signals were very low and in many cases of lower intensity than punctual auto-fluorescence signals seen in wild-type cells.

To determine whether the fluorescence signals truly originated from a knock-in at the desired genomic locus, several of the fluorescence positive clones of each knock-in experiment were tested for expression of the respective chimeric mRNA as described above. All five tested *CRY1* knock-in clones with clear nuclear fluorescence pattern were positive for the chimeric mRNA, while one tested clone with a different fluorescence pattern was not (**Supplementary Fig. S1C**). Out of seven tested fluorescence positive *PER2* knock-in clones, three were positive for the corresponding chimeric mRNA (**Supplementary Fig. S1C**). Sanger sequencing confirmed the identity of the PCR products.

Next, we examined the targeted genomic loci of these clones to check for precise integration of the transgenic DNA. To this end, we amplified the locus from genomic DNA of several clones using PCR primers that both bound outside the employed homology region to avoid any signal resulting from a randomly integrated vector (‘out-out-PCR’, **Supplementary Fig. S1D**). For most tested clones, we detected several PCR products: a major product corresponding to the size of the wild-type allele amplicon and minor products, including one with the expected size range of a knock-in allele amplicon, indicating a mono-allelic integration. For some tested clones, only the knock-in product was detected (e.g. one *CRY1*-mClover3 and one *PER2*-mScarlet-I clone), indicating knock-in at both alleles. Additional products are probably due to formation of heteroduplexes between wild-type and knock-in PCR products. Sanger sequencing of all tested knock-in alleles revealed exact matches with the predicted sequences for a successful knock-in, and confirmed a precise excision of the positive selection cassette. We also sequenced the second alleles of the mono-allelic integration clones. While the coding sequence was wild-type in two cases, Cas9 induced insertions or deletions for the other clones, resulting in alterations of up to 18 amino acids at the C-terminus. For one clone (PER2-mScarlet-I), a larger deletion (∼550 bp) was observed at the second allele (**Supplementary Fig. S1D**).

Finally, we wanted to exclude that the observed fluorescence signals originate from any other source than the PER2- or CRY1-fusion proteins (such as additional random FP integration). To this end, we transduced the knock-in clones with shRNA targeting *PER2* or *CRY1* mRNA and recorded fluorescence over 24 hours (**Fig. 1F, Supplementary Fig. S2**). In *CRY1* knock-in cells treated with shRNA against *CRY1* fluorescence signals were almost absent (with only auto-fluorescence visible), while cells left untreated or treated with shRNA against *PER2* showed robust nuclear fluorescence signals (red or green). Similarly, transient fluorescence signals in *PER2* knock-in cells appear even enhanced upon knockdown of *CRY1* (probably due to reduced repression of *PER2* transcription). In contrast, no signal exceeding auto-fluorescence was observed in these cells upon transduction with an anti-*PER2*-shRNA at any time point. Together, this strongly indicates that the observed fluorescence is due to proteins translated from the same mRNA as CRY1 or PER2 proteins, and thus fluorescence originates exclusively from the targeted fusion proteins.

Combining fluorescence, PCR and sequencing data, we observed that between 5 and 56 % (median 19 %) of the initially screened clones constitute successful knock-in clones (**Supplementary Fig. S1E**). For each knock-in (*PER2*-mClover3, *PER2*-mScarlet-I, *CRY1*-mClover3 and *CRY1*-mScarlet-I), we chose one clone for further experiments.

### Clock protein knock-in cells possess an intact circadian clock

Knocking-in fluorescent protein tags into *PER2* and *CRY1* genomic loci allows studying the endogenous clock protein’s dynamics in living cells if the knock-in does not affect the functionality of the molecular oscillator. To test this, we monitored *Bmal1*-promoter driven luciferase rhythms over several days in five selected clones (mono-allelic PER2-mClover3, mono- and bi-allelic PER2-mScarlet-I, bi-allelic CRY1-mClover3 and mono-allelic CRY1-mScarlet-I). All clones showed robust circadian oscillations with amplitudes and periods similar to wild-type U-2 OS cells (**Supplementary Fig. S3A-D**). This was expected for C-terminally tagged PER2, since homozygous *Per2*-luciferase or *Per2*-Venus knock-in mice show normal circadian locomotor behavior (*16, 21*). Adding a C-terminal tag to CRY1 could in principle lead to a hypomorphic allele with altered functionality of the corresponding fusion protein, which might go undetected in cells with only one allele carrying the knock-in and the other essentially being wild-type. This is unlikely, however, since (i) various C-terminal tags did not alter CRY1’s ability to repress CLOCK:BMAL1 transactivational activity (**Supplementary Fig. S3E**) and (ii) the CRY1-mClover3 clone with a bi-allelic knock-in showed essentially normal circadian dynamics (no period shortening as expected for hypomorphic alleles). In summary, the molecular clock in the tested knock-in cells was still functional and there was no indication that the fusion proteins represent non-functional variants.

### Protein dynamics of CRY1 and PER2

In the past, predominantly biochemistry experiments (with cell populations) suggested that both PERs and CRYs first accumulate in the cytoplasm, while their nuclear abundance shows circadian rhythms with peak levels in peripheral tissues at CT16-20 (*17, 18, 30*). To test whether and to which extent this is also true in individual living cells, we monitored fluorescence in synchronized knock-in cells at regular one-hour intervals over the course of three days.

In contrast to our expectations, the fluorescence of CRY1 fusion proteins was exclusively observed in the nucleus at any given time point (**Fig. 2A,B**), indicating that the majority of CRY1 is predominantly in this compartment irrespective of time of day. The nuclear signal intensity was well over background at all time-points in almost all cells and oscillated with a circadian period in the majority of cells. In contrast to CRY1, nuclear fluorescence signals of PER2 fusion proteins were more transient and detected for only 8-12 consecutive hours in an individual cell (**Fig. 2C,D**), resulting in circadian rhythms of PER2 nuclear signal consistent with previous reports from *Per2*-Venus knock-in mice (*16*). Prior to nuclear accumulation, a weak cytoplasmic PER2-FP signal was detectable in some cells, however, a reliable discrimination from auto-fluorescence and background signals was not possible (not shown).

**Figure 2:**
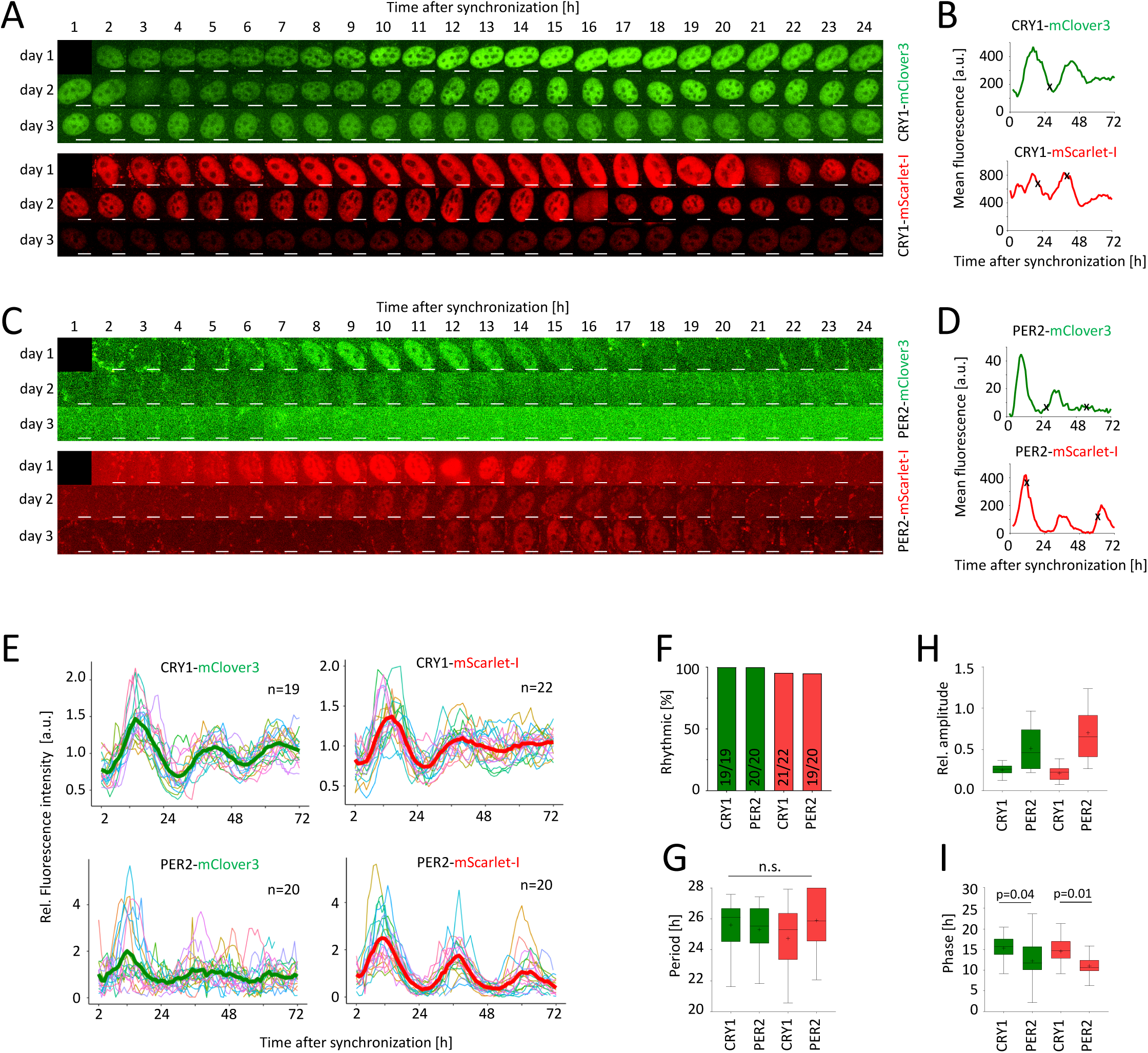
PER2- and CRY1-fusion protein oscillate in single knock-in cells. **(A)** and **(C)** Montage of fluorescence microscopy images of selected individual U-2 OS single knock-in cells’ nuclei over the course of 3 days after synchronization. **(B)** and **(D)** Mean of nuclear fluorescence intensity (background-subtracted) quantified from A and C. Cell division marked by (x). **(E)** Time series of normalized mean nuclear fluorescence in individual knock-in cells with average signal overlaid in bold. **(F)** Percentage of significantly rhythmic time series from E. **(G-I)** Extracted rhythm parameters of significantly rhythmic single cell time series from E. p-values: one-way ANOVA. Scale bar: 10 µm. Boxplots: Median with range, mean is marked with (+).

Together, we concluded that (i) the circadian clock was still intact in our knock-in clones, (ii) both CRY1 and PER2 fusion protein levels oscillate in a manner consistent with their well-established circadian regulation and (iii) PER2 and CRY1 expression dynamics can be monitored in the nuclei of single cells.

To exclude that the observed expression patterns are specific to U-2 OS cells, we also knocked-in dClover2 and mScarlet-I into the *PER2* and *CRY1* locus of the human colon epithelia cell line HCT-116 and monitored fluorescence over the course of two days (**Supplementary Fig. S4**). The spatiotemporal fluorescence patterns essentially recapitulated those seen in the U-2 OS cells: CRY1 was detectable almost exclusively in the nucleus over the whole circadian cycle, while nuclear PER2 fusion protein signal was detectable for less than 12 consecutive hours indicating that the observed dynamics of PER2 and CRY1 are similar in human cells.

### CRY1 is phase-delayed compared to PER2

To obtain a more quantitative picture of the spatiotemporal dynamics of PER2 and CRY1, we tracked nuclear fluorescence of ∼20 individual cells of each knock-in clone over three days (**Fig. 2E**). We used MetaCycle (*31*) to analyze the time series for the presence of circadian rhythms. Both CRY1-mScarlet-I and CRY1-mClover3 showed significant circadian rhythmicity of nuclear abundance in almost all cells with average periods of 24.7 ± 2.0 h and 25.6 ±1.6 h (mean ± SD), respectively (**Fig. 2F,G**). Circadian rhythms of nuclear fluorescence were also observed for the majority of *PER2*-mScarlet-I and *PER2*-mClover3 knock-in cells with average periods of 25.9 ± 2.0 h and 25.3 ± 1.7 h, respectively (**Fig. 2F,G**). The average relative amplitudes of PER2-fusion protein nuclear abundance were twice as high as these of CRY1-fusion proteins (**Fig. 2H**).

When comparing the average phases of PER2 and CRY1 protein rhythms in the nucleus, CRY1 fusion proteins appear to be phase-delayed relative to corresponding PER2 proteins by more than 3 h (**Fig. 2I**). However, at this stage it remained unclear whether this reflects phase differences in individual cells or resulted from clonal variation. We also observed a minor phase advance of the Scarlet-fusion proteins compared to the corresponding mClover3 fusion proteins of ∼1 h (**Fig. 2I**), which – in addition to clonal variation – possibly reflects differences in maturation time.

### Generation of double knock-in cells

To test whether delayed nuclear CRY1 accumulation is due to variability between individual cells or whether it is indeed a feature of the circadian oscillator, we generated knock-in cells expressing both CRY1 and PER2 as fluorescence tagged fusion proteins in different colors. To this end, we used PER2-mClover3 and PER2-mScarlet-I mono-allelic knock-in cells to generate double knock-in cells by integration of the complementary fluorophore into the CRY1 locus as described above (**Fig. 1A,B**). After positive and negative selection and CRE-mediated excision of the selection cassette, single clones were screened by microscopy. Again, high proportions of the inspected clones (12 out of 14 (86 %) for CRY1-mScarlet-I knock-in and 11 out of 19 (58 %) for CRY1-mClover3 knock-in) showed similar nuclear pattern as seen in the CRY1 single knock-in clones (**Fig. 3A,B, Supplementary Fig. S5A**). Three clones of each were selected, and RT-PCR and PCR/sequencing of the *CRY1* locus confirmed successful knock-in for all clones (**Supplementary Fig. S5B,C**). As expected, the spatial fluorescence patterns of PER2 and CRY1 fusion proteins substantially overlapped in the nucleus (**Fig. 3A**). Furthermore, circadian rhythms were still intact in those cells as confirmed by bioluminescence imaging using a *Bmal1*-luciferase reporter (**Supplementary Fig. S5D-G**).

**Figure 3:**
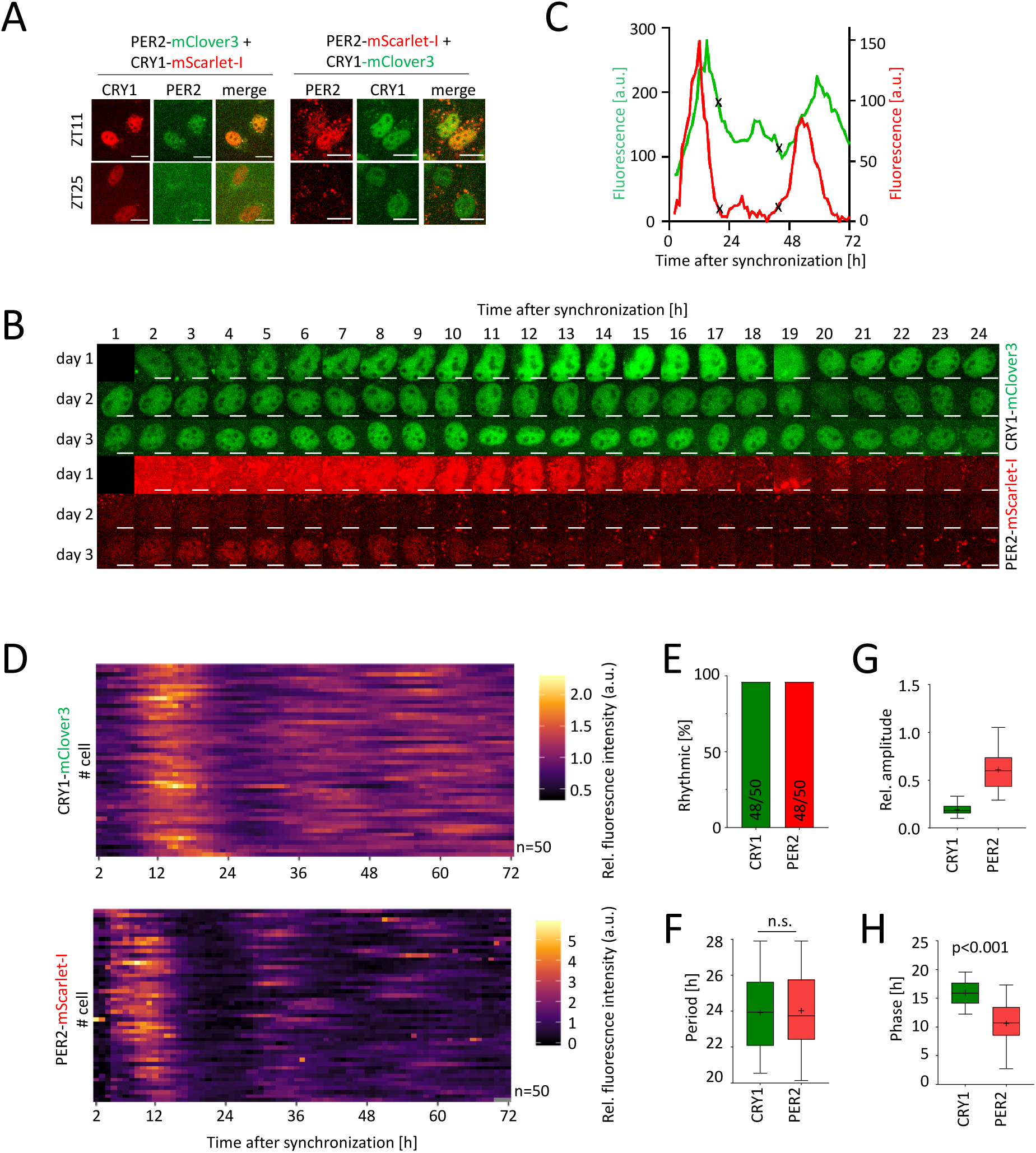
Simultaneous visualization of PER2- and CRY1-fusion protein oscillations in double knock-in cells. **(A)** PER2- and CRY1-fusion protein oscillation in individual double knock-in cells. Fluorescence images of selected double-knock in clones at different times after synchronization. **(B)** Montage of bicolor fluorescence microscopy images of an individual U-2 OS double-knock-in cell’s nucleus over the course of 3 days after synchronization. **(C)** Mean nuclear fluorescence intensity (background-subtracted) quantified from B. Cell division marked by (x). **(D)** Time series of normalized mean nuclear fluorescence in individual double knock-in cells. **(E)** Percentage of significantly rhythmic time series from E. **(F-H)** Extracted rhythm parameters of significantly rhythmic single cell time series from E. p-values: unpaired Student’s t-test. Boxplots: Median with range, mean is marked by (+). Scale bar: 10 µm.

### CRY1 is phase-delayed compared to PER2 also in individual cells

To quantify the temporal relationship of nuclear CRY1 and PER2 protein expression, we synchronized *CRY1*-mClover3/*PER2*-mScarlet-I double knock-in cells and determined nuclear fluorescence intensity of 50 individual cells over the course of 3 days (**Fig. 3B-D**). Again, almost all cells (>95 %) displayed significant rhythmicity of CRY1 and PER2 levels with average periods of 24.0 ± 2.3 h for both proteins and a ∼3-fold higher relative amplitude of PER2 rhythms (**Fig. 3E-G**). As seen in the single knock-in cells, mean CRY1-mClover3 nuclear fluorescence signal followed that of PER2-mScarlet with a delay of ∼5 h (**Fig. 3C,H**). In individual cells, the median phase difference between PER2 and CRY1 nuclear abundance rhythms was 5.4 h (**Fig. 4A**). To test whether this phase difference is dominated by differential responses of PER2 and CRY1 expression to dexamethasone synchronization, we reanalyzed the time series starting from 26 h post treatment, thus omitting data from the first circadian cycle. Again, the phase of CRY1 rhythmicity was still delayed by 4.9 h compared to PER2 rhythms (**Fig. 4B**) indicating that the observed delay was not due to acute dexamethasone effects.

**Figure 4:**
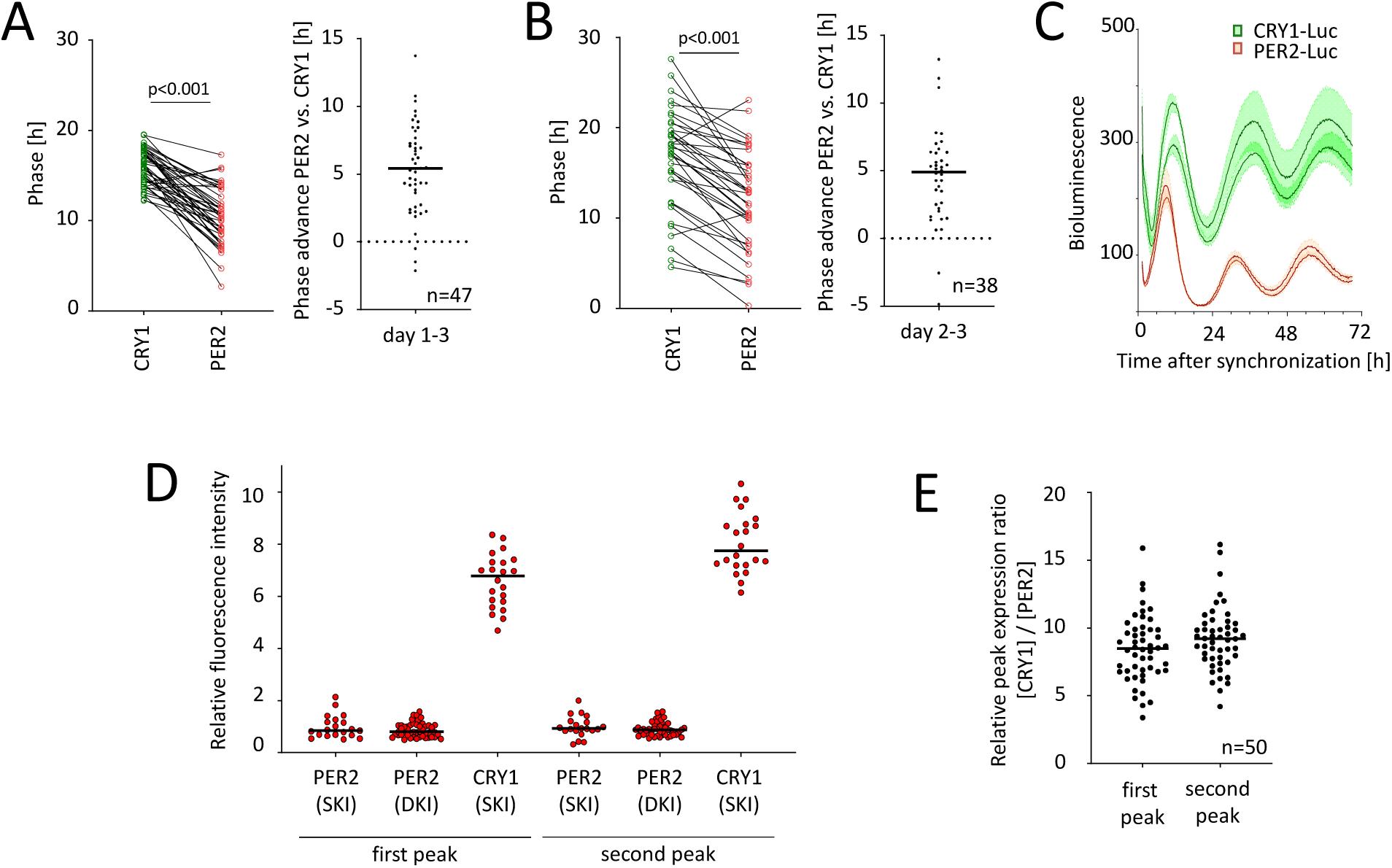
Nuclear CRY1 peaks much later and is much more abundant than PER2. **(A-B)** Analysis of phase difference between CRY1 and PER2 nuclear accumulation in individual double knock in cells. Phases were calculated either including (A) or excluding (B) the first 24 hours of the three days’ time series. p-values: paired Student’s t-test. **(C)** Live-cell bioluminescence recordings of knock-in cells expressing either CRY1-luciferase or PER2-luciferase fusion proteins. Depicted are mean ±SD of 3 individual traces from 2 clones of each knock-in. Representative data from 2 experiments. **(D)** Relative nuclear peak intensity of fluorescence in cells expressing either PER2- or CRY1-mScarlet fusion protein. **(E)** Ratio of normalized expression of CRY1-mClover3 versus PER2-mScarlet-I in individual cells expressing both fusion proteins. Lines in scatter plots depict median. SKI: single knock-in. DKI: double knock-in.

To investigate whether the PER2-CRY1 phase difference is specific for *nuclear* accumulation or whether it is a feature of their whole-cell expression dynamics, we knocked-in firefly luciferase into the *CRY1* or *PER2* loci of U-2 OS cells and recorded luminescence of three individual clones over the course of three days. (Reliable quantification of whole-cell fluorescence signals was impossible, because of the rather low fluorescence and high auto-fluorescence signals in the cytoplasm of FP-reporter cells). As observed for fluorescence fusion proteins, the phase of CRY1-LUC expression rhythms was delayed by ∼5 h compared to that of PER2-LUC (**Fig. 4C**).

### CRY1 is much more abundant than PER2 in the nucleus of U-2 OS cells

Mass spectrometry data from mouse liver suggested that CRY1 protein peak levels are higher than those of PER2 (*28*). Since we have tagged CRY1 and PER2 with the same fluorescent proteins, we could quantitatively compare the expression intensities of both proteins in U-2 OS cell nuclei. To this end, we quantified peak fluorescence levels, from bi-allelic PER2-mScarlet-I knock-in cells, from mono-allelic CRY1-mScarlet-I knock-in cells (we do not have bi-allelic knock-in cells yet), and from double knock-in cells expressing PER2-mScarlet-I from one allele (and also CRY1-mClover3 from one allele). Assuming that the distribution of peak expression levels are roughly similar between the different knock-in cell lines, and that the signal from two alleles would be additive, we estimated that average peak expression of CRY1 is more than six times higher than that of PER2 (**Fig. 4D**). Comparable ratios were obtained by comparing peak expression of mClover3 fusion proteins (not shown). In a similar manner, CRY1-luciferase knock-in cells gave rise to much higher signals in comparison to PER2-luciferase knock-in cells on a population level, indicating that these abundance level differences are not confined to the nucleus (**Fig. 4C**).

To compare peak intensities between PER2 and CRY1 on single cell level, we took advantage of the fact that we have different cell lines expressing CRY1 coupled to either mScarlet-I or mClover3. Again assuming that the distribution of CRY1 peak expression levels is roughly similar among the different knock-in cell lines, we compared the signal intensities of CRY1-mScarlet-I and CRY1-mClover3 at the peak of circadian expression. This allowed us to estimate the relative amount of PER2-mScarlet-I and CRY1-mClover3 in individual double-knock-in cells. Similar to the population level, the peak amount of CRY1 surpassed that of PER2 in the same cells by 8.5 ± 2.4 fold for the first peak and by 9.2 ± 2.3 fold for second peak (**Fig. 4E**), suggesting that CRY1 protein is present at much higher levels than PER2 protein in the nucleus of U-2 OS cells.

## DISCUSSION

The current model of the mammalian circadian oscillator is predominantly based on data from genetics and biochemistry experiments that have been accumulated over more than 20 years. In addition, luciferase reporter technology substantially advanced our knowledge about the dynamics of circadian rhythms in live cells. The cell biology of circadian clocks, however, is still in its infancy mainly due to the lack of suitable reporter technologies that allow the (simultaneous) spatiotemporal quantification of individual clock proteins in living single cells. This is likely due to the fact that even almost a decade after the CRISPR revolution, the generation of knock-in cell lines is still not a standard technique, but a time consuming endeavor with uncertain success. Here, we enrich the existing toolbox by an efficient selection strategy that is particularly useful for lowly or transiently expressed genes, such as those coding for circadian clock proteins. The selection process is independent of target gene expression and only leaves a single loxP site in the genome. For achieving high knock-in efficiencies, our approach can be combined with complementary techniques, such as the use of CRISPR/Cpf1 or cell cycle synchronization (*32, 33*).

The resulting fluorescent knock-in cell lines enabled us to visualize endogenously expressed PER2 and CRY1 proteins – two canonical circadian repressors – for the first time in single human live cells. We are confident that the fusion proteins are functional, because cells with a bi-allelic knock-in of PER2- or CRY1-fusion proteins display normal circadian rhythms, in contrast to *Per2* or *Cry1* knockout cells which are hardly rhythmic (*34, 35*). This is further supported by reports that homozygous *Per2*-Venus and *Per2*-Luc knock-in mice display normal circadian rhythms, and rhythmically expressed CRY1-EGFP can rescue rhythmic behavior in otherwise arrhythmic *Cry1/2* double knock-out mice (*16, 36*). Therefore, these knock-in cells should constitute reliable tools to study the spatiotemporal dynamics of both proteins within a widely used model of the human circadian oscillator on the single cell level.

Circadian oscillations occur because of a critical delay in the auto-regulatory negative feedback of PER and CRY proteins on their own transcription. Delayed nuclear accumulation of negative regulators has been discussed to be one mechanism in this context. Indeed, in mammals, nuclear levels of PER and CRY proteins seem to mutually depend on the presence of the other protein family members (*12*). In *Drosophila*, the analogs of PER and CRY, dPER and dTIM, are reported to first accumulate in the cytoplasm for several hours and then translocate into the nucleus in a switch-like event (*37*). In mouse liver, biochemistry data suggest that CRY and PER proteins almost exclusively coexist in huge cytosolic and nuclear complexes suggesting common regulation (*17*). In contrast, several biochemical studies reported the presence of nuclear CRY1 protein over the whole circadian cycle in human cell lines and mouse liver (*12, 18, 38*), but it remained unclear whether this was due to sampling populations of unsynchronized cells.

Using our new reporter cells, we found three interesting features of circadian clock protein dynamics in human cells that raise questions regarding the current model of the circadian oscillator. Firstly, CRY1 is in the nucleus at all circadian phases in U-2 OS cells (and also in the colon epithelial cell line HCT-116) with no detectable CRY1 in the cytoplasm. Thus, the majority of translated CRY1 protein seems to immediately enter the nucleus without any circadian gating. Similarly, we did not observe major cytoplasmic accumulation of PER2 protein prior to its nuclear appearance consistent with data from cells from the *Per2*-Venus (*16*). Secondly, nuclear PER2-levels peak on average ∼5 h before CRY1-levels in single cells consistent with data from population sampling of murine cell nuclei (*12, 30, 39*). This delay is also present on whole-cell protein level (**Fig. 4C** and (*28*)) indicating that in single cells circadian nuclear accumulation of CRY1 and PER2 mainly reflects the circadian expression of those proteins rather than being the consequence of circadian gating in nuclear appearance. Thirdly, quantification of fluorescence signal from fusion proteins indicated that CRY1 peak levels exceed those of PER2 by a factor of ∼6-10 in U-2 OS cells. This is a much larger difference than that seen in mouse liver, where CRY1 peak expression level was reported to be only twice as high as that of PER2 (*28*).

Together, these data raise the following questions: (i) Does CRY1 nuclear accumulation really directly depend on the presence of PER proteins as previously suggested (*12*)? Since CRY1 is mainly nuclear regardless of PER2 expression phase (peak or trough), it peaks when PER2 levels are already declining, and it is 6-10-fold more abundant, at least PER2 levels seem not to be a limiting factor for CRY1 nuclear entry. Since biochemistry data indicate the existence of cytoplasmic complexes that contain CRY proteins but not PER2 in mouse liver (*17*), it is possible that other PER protein family members (PER1 and/or PER3) act as CRY1 carriers for nuclear entry. Our knock-in technology should now allow the efficient generation of PER1 and PER3 reporter cells to study this issue. It is also conceivable that a single PER2 protein may be able to shuttle multiple CRY proteins, either at a time or in a sequential manner. The described dynamic shuttling of PER2-Venus protein in and out of the nucleus supports the latter (*16*). (ii) Is CRY1 in the nucleus always almost exclusively present in a large negative feedback complex? Biochemistry data with murine liver lysates indicate that the majority of CRY1 protein indeed is present in a ∼1.9-MDa negative feedback complex, which also includes CLOCK-BMAL1 and virtually all of the PER and CRY proteins as well as CK1δ (*17*). Only a small minority of nuclear CRY1 was found as monomer. Although the stoichiometry of clock proteins within the murine negative feedback complex has not yet been worked out, this seems to be different in U-2 OS cells. The delayed phase of nuclear CRY1 abundance compared to PER2, the fact that PER2 and CRY1 directly interact in a 1:1 ratio (*9*) and, more importantly, the much higher protein abundance at all circadian phases point to a much higher degree of CRY1 proteins not being in a complex with PER2 in human cells. It will be interesting to investigate the PER2-independent targets of CRY1, which may include nuclear receptors (*40*).

In summary, we present a novel CRISPR-based knock-in strategy that allows the efficient generation of reporter cells even for genes with low or variable expression levels, such as circadian clock genes. We created PER2 and CRY1 single and double knock-in reporter cells with clock proteins tagged to fluorescence proteins or luciferase. These, for the first time, enabled the visualization of PER2 and CRY1 protein dynamics in live single cells. Although individual cellular oscillators display a pretty large degree of intercellular variability with respect to dynamic parameters, we discovered several features of the (human) circadian oscillator that are not easily consistent with the canonical circadian oscillator model. Future studies are required to evaluate whether these differences are due to the species, the tissue, the detection methods or other unknown factors. We anticipate that the generation of additional single, double or triple knock-in cells for circadian clock proteins will greatly advance our understanding about the cell biology of circadian clocks. Our study is the first step.

## MATERIALS AND METHODS

### Cell lines

U-2 OS (human, ATCC HTB-96) and HCT-116 (human, ATCC CCL-247) cells were cultured in DMEM supplemented with 10 % FBS, 25 mM HEPES and penicillin/streptomycin at 37 °C and 5 % CO_2_. For long-term imaging, cells were cultured in FluorBrite (GIBCO) medium supplemented with 2 % FBS, 1x GlutaMax and penicillin/streptomycin from two days prior to imaging.

### Plasmids

The core sequence including positive selection cassette, LoxP sites, Frt-sites, His-Flag-tag, poly(A)-sites and multiple cloning sites for insertion of homology arms was synthesized by commercial supplier (BaseClear) and cloned into pUC19 backbone. A negative selection cassette with human thymidine kinase was retrieved from Addgene #21911 (*41*). The hTK was exchanged for hCD4, amplified from the pMSCV-IRES-hCD4plasmid (Addgene #35712 (*42*)). mClover3 was subcloned from Addgene #74252 (*26*), mScarlet was subcloned from Addgene #98839 (*43*). HaloTag and SNAP-Tag sequences are listed in **Supplementary Table S1** and were subcloned from expression vectors. The pCAG-i53bp expression plasmid was a gift from Ralf Kuhn and is derived from Addgene #74939 (*29*). SV40-NLS CRE recombinase was a gift from Christoph Harms and was subcloned into pLenti6 backbone. CRY1 fusion proteins also included translation of the LoxP site C-terminal to the fluorophore to avoid nonsense-mediated decay. See **Supplementary Table S1** for DNA sequence information.

Single guide RNAs (**Supplementary Table S2**) were designed to cut near the STOP codon using CRISPOR (*44*), and corresponding DNA oligos were ligated into pCRISPR-Lenti-v2 (Addgene #52961 (*45*)) as described. To test efficiency of guides, cells were transduced with lentiviruses harboring the Cas9/sgRNA expression plasmid, puromycin resistant cells’ gDNA was isolated, the respective region amplified by PCR and sequenced. Efficiency was assessed using TIDE assay (*46*).

pGIPZ clones V2LHS_172866 (CRY1) V2LHS_52938 (PER2) (**Supplementary Table S3**) were purchased from Open Biosystems (GE Healthcare) and the tGFP was mutated to abrogate fluorescence. The 0.9 kb *Bmal1*-promoter driven luciferase reporter construct is described in (*47*).

### Transfection

For knock-in experiments, 10^6^ cells were harvested by trypsinization and transfected with each 2 µg of i53bp, donor vector, and pCRSIPR-Lenti-V2 by electroporation using the NEON system (Thermo Fisher, buffer N, U-2 OS: 4 pulses, 10 ms, 1230 V, HCT-116: 2 pulses, 30 ms, 1130 V). For *PER2*-editing, a mixture of three sgRNA sequences was used in equimolar ratios. After electroporation, cells were seeded into antibiotic-free DMEM and cultured for 24 hours before selection. Transient transfections of CRE recombinase were performed using 1 µL Lipofectamin 2000 and 200 ng CRE expression plasmid in a 48-well plate format.

### Virus production and transduction of cells using lentivirus

HEK293-T cells were transiently transfected in a T75 flask with 8.6 µg lentiviral expression plasmid, 6 µg psPAX2 and 3.6 µg pMD2G (gift from the Trono lab, Addgene #12259 and #12260) packaging plasmid using CalPhos Mammalian Transfection Kit (Takara). Next day, culture medium was replaced by 12.5 mL complete culture medium, and lentiviral supernatant was collected after 24 and 48 h. Combined supernatant was passed through a 0.45 µm filter (Filtropur S 0.45) and either used directly or stored in aliquots at −80 °C. For transduction, cells were seeded into lentivirus containing supernatant supplemented with 8 µg/mL protamine sulphate. Next day, lentivirus containing supernatant was aspired and cells were cultured in complete culture medium for further 24 h before antibiotic selection of transduced cells.

### Antibiotic selection

To select for transfected or transduced cells, cells were grown sub-confluently in blasticidin (10 µg/ml) containing medium for > 3 days or in puromycin (10 µg/ml) containing medium for > 1 day, until non-transfected control cells died.

### FACS sorting

Cells were sorted on a FACS AriaII (BD). For staining of surface hCD4 for negative selection, 2×10^6^ cells were trypsinized, washed with 0.5 % BSA/PBS and incubated with 200 µL of a 1:50 dilution of hCD4-BV711 (OKT4, BioLegend, UK) for 30 min. Cells were washed twice with BSA/PBS. Excitation: 405 nm. Emission filter: 525LP-525/50 (CFP), 685LP-710/50 (BV711).

### Nucleic acid isolation and PCR

Genomic DNA was extracted with direct PCR Lysis Reagent Cell (VWR). RNA isolation was performed using the AMBION PureLink RNA Mini kit (Themo Fisher) according to the manufacturer’s instruction, including an on-column DNase digest. RNA was reversely transcribed using gene specific primer and two-step protocol. PCR amplifications were performed using Phusion polymerase (New England Biolabs), products were analyzed by agarose gel electrophoresis and detected using RedSafe/UV light. Primer sequences are listed in **Supplementary Table S4**. Of notice, additional low mobility bands were observed from PCR of one-allelic knock-in clones (**Supplementary Figure S1D** and **S5C**). Sanger sequencing of one of those bands revealed a mixture of wild-type and knock-in sequence. This and the fact that we did not observe low mobility bands in PCRs from bi-allelic knock-in clones made us conclude that they represent slowly migrating heteroduplexes and not additional PCR products.

### Bioluminescence recording of circadian oscillation

Luciferase knock-in cells or fluorescent knock-in cells transduced with an m*Bmal1* promoter driven luciferase reporter plasmid were seeded to reach confluence. To synchronize the circadian rhythms, cells were either washed twice with cold PBS for 2 min (fluorescent knock-in cells), or treated with 1 µM dexamethasone for 30 min followed by washing with warm PBS (luciferase knock-in cells). Cells were then incubated in DMEM without phenol-red supplemented with 250 µM D-luciferin, and dishes were sealed using parafilm. Bioluminescence was recorded in a 96-well plate luminometer (TopCount, Perkin Elmer) or in a LumiCycle (Actimetrics). Raw data were detrended by dividing by the 24-h running average. Periods, phases and mean bioluminescence signal were estimated by fitting the cosine wave function using the ChronoStar software as described (*48*).

### Microscopy

For Microscopy, cells were seeded on glass bottom #1.5H µ-slides (IBIDI, Germany) or glass bottom #1.5H-N 96 well plates (Cellvis, USA). Imaging was performed on a Nikon Widefield Ti2 equipped with a sCMOS, PCO.edge camera and a live-cell incubator. Image acquisition was done in Fluorbrite medium (GIBCO) supplemented with 2 % FBS, 1:100 PenStrep, and 1x GlutaMax at 37 °C and 5 % CO_2_. The following light sources (LEDs) and emission filters were used for the different channels: CFP (cerulean): excitation 438/29, emission 473/24 nm; GFP (dClover2): excitation 475/28 nm, emission 520/26 nm; YFP (mClover3, dClover2): excitation 511/16 nm, emission 540/30 nm; RFP (mScarlet-I) excitation 555/28 nm, emission 642/80 nm. Objectives: 40x ApoFluor, NA 0.95, WD 250 µm; 20x Plan Apo, NA 0.8, WD 1 mm. Typically, illumination time for CFP was 500 ms and 2 s for all other channels. For time course experiments, cells were synchronized by addition of 1 µM dexamethasone for 30 min followed by washing with warm PBS, and imaging started 2 h after synchronization with a regular imaging interval of 1 h.

### Fluorescence data analysis

Fluorescence data were extracted using ImageJ. Nuclei were manually marked using either the respective fluorescence channel (CRY1) or phase contrast (PER2), and mean fluorescence intensities were extracted. Individual background fluorescence for every cell at every time point was determined by quantifying the same area of the imaging field from a cell-free image frame of the same experiment. Mean background signal was subtracted. Linear trends were detected by linear regression analysis of all cells from one imaging frame and also eliminated by subtraction. Since trough PER2-fluorescence levels were not surpassing background noise, we set the lowest intensity of each time series to 0. To eliminate cell division outliers, fluorescence values at cell division events were imputed by averaging across neighboring time points.

The data were exported, processed and analyzed in R (version 3.6). Rhythmicity of single cell time-series was evaluated in with meta2d from the MetaCycle R package (version 1.2 (*31*)), with minper=20, maxper=28, and cycMethod=c (“ARS”, “JTK”, “LS”), thus incorporating the ARSER, JTK_CYCLE and Lomb-Scargle algorithms (*49–51*). Time series that did not pass the rhythmicity test (Benjamini-Hochberg FDR>0.05) were excluded from further analysis. Data were normalized for **Fig. 2E** and **Fig. 3D** by dividing intensities by mean intensities of the respective time series. Plots were drawn using ggplot2 v3.2. To facilitate comparison of phases, extracted phase was divided by extracted period (mean period for double knock-in cells) and multiplied by 24. When comparing phases of double knock-in cells, we chose the direction (advance or delay) according to which absolute resulting phase difference was lower than 10, or, when in rare cases above 10, closer to the mean of all other data.

### Semi-quantitative analysis of fluorescence signals

To confidently compare fluorescence data for the different fusion proteins with the same fluorophore entity, we made the following assumptions and analyzed and normalized the data accordingly: (i) Because PER2 trough expression levels were not distinguishable from background, we compared peak expression levels that can be quantified with higher confidence. (ii) As we assumed that the distribution of peak protein expression is similar for different clones, we extracted peak fluorescence values from each background-subtracted time series in a time window of either the first or the second day of recording and compared the means. (iii) We assumed that the expression of PER2 protein from two alleles would be twice the expression from one allele. Comparing the first peaks of mono-and bi-allelic PER2-mScarlet-I expression, we added a constant (28.7 a.u.) to all peak values in order to fulfill this assumption best for the lower 60 % of all peaks (smallest squares of linear fit). Conceptually, this constant can be regarded as the lower limit of mScarlet-detection. This correction reduces the underestimation of fusion protein expression from low signals. For comparison, peak values from bi-allelic knock-in were divided by two, and all data were normalized to the mean of PER2-mScarlet-I peak expression of single knock-in cells (**Fig. 4C**).

To be able to estimate protein ratios in single cells, in which PER2 and CRY1 are fused to different fluorophores, we first had to determine the relative signal intensities of mScarlet-I and mClover3 resulting from a similar amount of fusion proteins in our experiment. We compared mean peak intensities (1^st^ and 2^nd^ peak independently) of CRY1-mClover3 (divided by two because of the bi-allelic knock-in) with those of CRY1-mScarlet-I from single-knock-in or double knock-in cells. We estimated that in average, mScarlet-I intensities surpass mClover3 intensities resulting from a supposedly similar amount of fluorophores by a factor of 4.3 under the used experimental settings. By correcting for this, and for the detection limit of PER2 (see above), the ratio of CRY1-mClover to PER2-mScarlet was determined by:

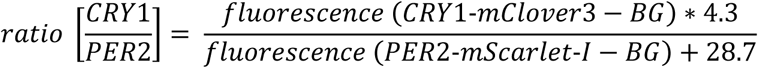

## ACKNOWLEDGEMENTS

We thank all current and former members of the Kramer and Herzel labs for technical and intellectual assistance. We would like to acknowledge the assistance of the FCCF at the Deutsches Rheuma-Forschungszentrum. Furthermore, we thank the Advanced Medical Bioimaging Core Facility (AMBIO) of the Charité for support in acquisition of the imaging data. We thank Mir-Farzin Mashreghi, Joachim Fuchs, Christoph Harms, Steve Brown and Michela Di Virgilio for providing material, and Janina Gabriel for assistance on data analysis. This work was funded by the Deutsche Forschungsgemeinschaft (DFG, German Research Foundation) - Project Number 278001972 - TRR 186.

## REFERENCES

1. N. Gekakis et al., Role of the CLOCK protein in the mammalian circadian mechanism. Science. 280, 1564–9 (1998).

2. K. Kume et al., mCRY1 and mCRY2 are essential components of the negative limb of the circadian clock feedback loop. Cell. 98, 193–205 (1999).

3. M. H. Vitaterna et al., Differential regulation of mammalian period genes and circadian rhythmicity by cryptochromes 1 and 2. Proc. Natl. Acad. Sci. U. S. A. 96, 12114–9 (1999).

4. S. H. Yoo et al., Competing E3 ubiquitin ligases govern circadian periodicity by degradation of CRY in nucleus and cytoplasm. Cell. 152, 1091–1105 (2013).

5. Y. Isojima et al., CKIepsilon/delta-dependent phosphorylation is a temperature-insensitive, period-determining process in the mammalian circadian clock. Proc. Natl. Acad. Sci. U. S. A. 106, 15744–9 (2009).

6. E. J. Eide et al., Control of mammalian circadian rhythm by CKIepsilon-regulated proteasome-mediated PER2 degradation. Mol. Cell. Biol. 25, 2795–807 (2005).

7. B. Zheng et al., Nonredundant Roles of the mPer1 and mPer2 Genes in the Mammalian Circadian Clock. Cell. 105, 683–694 (2001).

8. G. T. van der Horst et al., Mammalian Cry1 and Cry2 are essential for maintenance of circadian rhythms. Nature. 398, 627–30 (1999).

9. I. Schmalen et al., Interaction of Circadian Clock Proteins CRY1 and PER2 Is Modulated by Zinc Binding and Disulfide Bond Formation. Cell. 157, 1203–1215 (2014).

10. R. Ye, C. P. Selby, N. Ozturk, Y. Annayev, A. Sancar, Biochemical analysis of the canonical model for the mammalian circadian clock. J. Biol. Chem. 286, 25891–25902 (2011).

11. S. N. Nangle et al., Molecular assembly of the period-cryptochrome circadian transcriptional repressor complex. Elife. 3, 1–14 (2014).

12. R. Ye et al., Dual modes of CLOCK:BMAL1 inhibition mediated by Cryptochrome and Period proteins in the mammalian circadian clock. Genes Dev. 28, 1989–98 (2014).

13. R. Ollinger et al., Dynamics of the circadian clock protein PERIOD2 in living cells. J. Cell Sci. 127, 4322–4328 (2014).

14. P. Meyer, L. Saez, M. W. Young, PER-TIM interactions in living Drosophila cells: an interval timer for the circadian clock. Science. 311, 226–9 (2006).

15. O. Tataroğlu, T. Schafmeier, Of switches and hourglasses: regulation of subcellular traffic in circadian clocks by phosphorylation. EMBO Rep. 11, 927–35 (2010).

16. N. J. Smyllie et al., Visualizing and Quantifying Intracellular Behavior and Abundance of the Core Circadian Clock Protein PERIOD2. Curr. Biol. 26, 1880–1886 (2016).

17. R. P. Aryal et al., Macromolecular Assemblies of the Mammalian Circadian Clock. Mol. Cell. 67, 770-782.e6 (2017).

18. Y. Lee, A. Reum Jang, L. J. Francey, A. Sehgal, J. B. Hogenesch, KPNB1 mediates PER/CRY nuclear translocation and circadian clock function. Elife. 4, 1–16 (2015).

19. H. R. Ueda et al., System-level identification of transcriptional circuits underlying mammalian circadian clocks. Nat. Genet. 37, 187–192 (2005).

20. R. Chen et al., Rhythmic PER Abundance Defines a Critical Nodal Point for Negative Feedback within the Circadian Clock Mechanism. MOLCEL. 36, 417–430 (2009).

21. S.-H. Yoo et al., PERIOD2::LUCIFERASE real-time reporting of circadian dynamics reveals persistent circadian oscillations in mouse peripheral tissues. Proc. Natl. Acad. Sci. U. S. A. 101, 5339–5346 (2004).

22. L. S. Mure et al., Diurnal transcriptome atlas of a primate across major neural and peripheral tissues. Science (80-.). 0318, 1–16 (2018).

23. T. Gaj, C. a Gersbach, C. F. Barbas, ZFN, TALEN, and CRISPR/Cas-based methods for genome engineering. Trends Biotechnol. 31, 397–405 (2013).

24. J.-S. Kim, Genome editing comes of age. Nat. Protoc. 11, 1573–1578 (2016).

25. V. Singh, D. Braddick, P. K. Dhar, Exploring the potential of genome editing CRISPR-Cas9 technology. Gene. 599, 1–18 (2017).

26. B. T. Bajar et al., Improving brightness and photostability of green and red fluorescent proteins for live cell imaging and FRET reporting. Sci. Rep. 6, 20889 (2016).

27. D. S. Bindels et al., MScarlet: A bright monomeric red fluorescent protein for cellular imaging. Nat. Methods. 14, 53–56 (2016).

28. R. Narumi et al., Mass spectrometry-based absolute quantification reveals rhythmic variation of mouse circadian clock proteins. Proc. Natl. Acad. Sci. U. S. A. 113, E3461–7 (2016).

29. M. D. Canny et al., Inhibition of 53BP1 favors homology-dependent DNA repair and increases CRISPR-Cas9 genome-editing efficiency. Nat. Biotechnol. 36, 95–102 (2018).

30. J. Wang et al., Nuclear Proteomics Uncovers Diurnal Regulatory Landscapes in Mouse Liver. Cell Metab. 25, 102–117 (2017).

31. G. Wu, R. C. Anafi, M. E. Hughes, K. Kornacker, J. B. Hogenesch, MetaCycle: an integrated R package to evaluate periodicity in large scale data. Bioinformatics. 32, 3351–3353 (2016).

32. S. Lin, B. T. Staahl, R. K. Alla, J. A. Doudna, Enhanced homology-directed human genome engineering by controlled timing of CRISPR/Cas9 delivery. Elife. 3, e04766 (2014).

33. B. Zetsche et al., Cpf1 is a single RNA-guided endonuclease of a class 2 CRISPR-Cas system. Cell. 163, 759–71 (2015).

34. T. Börding, A. N. Abdo, B. Maier, C. Gabriel, A. Kramer, Generation of human CrY1 and Cry2 knockout cells using duplex CRISPR/Cas9 technology. Front. Physiol. 10, 1–9 (2019).

35. A. C. Liu et al., Intercellular Coupling Confers Robustness against Mutations in the SCN Circadian Clock Network. Cell. 129, 605–616 (2007).

36. M. D. Edwards, M. Brancaccio, J. E. Chesham, E. S. Maywood, M. H. Hastings, Rhythmic expression of cryptochrome induces the circadian clock of arrhythmic suprachiasmatic nuclei through arginine vasopressin signaling. Proc. Natl. Acad. Sci. 113, 201519044 (2016).

37. P. Meyer, L. Saez, M. W. Young, PER-TIM interactions in living Drosophila cells: an interval timer for the circadian clock. Science. 311, 226–9 (2006).

38. A. Hirano, D. Braas, Y.-H. Fu, L. J. Ptáček, FAD Regulates CRYPTOCHROME Protein Stability and Circadian Clock in Mice. Cell Rep. 19, 255–266 (2017).

39. N. Koike et al., Transcriptional Architecture and Chromatin Landscape of the Core Circadian Clock in Mammals. Science (80-.). 338, 349–354 (2012).

40. A. Kriebs et al., Circadian repressors CRY1 and CRY2 broadly interact with nuclear receptors and modulate transcriptional activity. Proc. Natl. Acad. Sci., 201704955 (2017).

41. R. A. Dewey et al., Chronic brain inflammation and persistent herpes simplex virus 1 thymidine kinase expression in survivors of syngeneic glioma treated by adenovirus-mediated gene therapy: implications for clinical trials. Nat. Med. 5, 1256–63 (1999).

42. E. L. Artinger et al., An MLL-dependent network sustains hematopoiesis. Proc. Natl. Acad. Sci. U. S. A. 110, 12000–5 (2013).

43. M. Mastop et al., Characterization of a spectrally diverse set of fluorescent proteins as FRET acceptors for mTurquoise2. Sci. Rep. 7, 11999 (2017).

44. M. Haeussler et al., Evaluation of off-target and on-target scoring algorithms and integration into the guide RNA selection tool CRISPOR. Genome Biol. 17, 148 (2016).

45. N. E. Sanjana, O. Shalem, F. Zhang, Improved vectors and genome-wide libraries for CRISPR screening. Nat. Methods. 11, 783–784 (2014).

46. E. K. Brinkman, T. Chen, M. Amendola, B. van Steensel, Easy quantitative assessment of genome editing by sequence trace decomposition. Nucleic Acids Res. 42, e168–e168 (2014).

47. B. Maier et al., A large-scale functional RNAi screen reveals a role for CK2 in the mammalian circadian clock. Genes Dev. 23, 708–18 (2009).

48. B. Maier, S. Lorenzen, A.-M. Finger, H.-P. Herzel, A. Kramer, in Circadian Clocks: Methods and Protocols, S. Brown, Ed. (Springer, 2020).

49. M. E. Hughes, J. B. Hogenesch, K. Kornacker, JTK_CYCLE: an efficient nonparametric algorithm for detecting rhythmic components in genome-scale data sets. J. Biol. Rhythms. 25, 372–80 (2010).

50. E. F. Glynn, J. Chen, A. R. Mushegian, Detecting periodic patterns in unevenly spaced gene expression time series using Lomb-Scargle periodograms. Bioinformatics. 22, 310–6 (2006).

51. R. Yang, Z. Su, Analyzing circadian expression data by harmonic regression based on autoregressive spectral estimation. Bioinformatics. 26, i168–i174 (2010).

