## Supplementary Tables and Figures for "Live-cell imaging of circadian clock protein dynamics in CRISPR-generated knock-in cells"

**Supplementary Table S1: DNA sequences**

| Part | Sequence |
|---|---|
| **His-Flag-Tag (HF-tag)** | CACCATCACCATCACCATGGTAGCGGTGACTACAAAGACGATGACGACAAG |
| **hCD4 extracellular domain** | ATGAACCGGGGAGTCCCTTTTAGGCACTTGCTTCTGGTGCTGCAACTGGCGCTCCTCCCA<br>GCAGCCACTCAGGGAAAGAAAGTGGTGCTGGGCAAAAAAGGGGATACAGTGGAACTGACC<br>TGTACAGCTTCCCAGAAGAAGAGCATACAATTCCACTGGAAAAACTCCAACCAGATAAAG<br>ATTCTGGGAAATCAGGGCTCCTTCTTAACTAAAGGTCCATCCAAGCTGAATGATCGCGCT<br>GACTCAAGAAGAAGCCTTTGGGACCAAGGAAACTTCCCCCTGATCATCAAGAATCTTAAG<br>ATAGAAGACTCAGATACTTACATCTGTGAAGTGGAGGACCAGAAGGAGGAGGTGCAATTG<br>CTAGTGTTCGGATTGACTGCCAACTCTGACACCCACCTGCTTCAGGGGCAGAGCCTGACC<br>CTGACCTTGGAGAGCCCCCCTGGTAGTAGCCCCTCAGTGCAATGTAGGAGTCCAAGGGGT<br>AAAAACATACAGGGGGGGAAGACCCTCTCCGTGTCTCAGCTGGAGCTCCAGGATAGTGGC<br>ACCTGGACATGCACTGTCTTGCAGAACCAGAAGAAGGTGGAGTTCAAAATAGACATCGTG<br>GTGCTAGCTTTCCAGAAGGCCTCCAGCATAGTCTATAAGAAAGAGGGGGAACAGGTGGAG<br>TTCTCCTTCCCACTCGCCTTTACAGTTGAAAAGCTGACGGGCAGTGGCGAGCTGTGGTGG<br>CAGGCGGAGAGGGCTTCCTCCTCCAAGTCTTGGATCACCTTTGACCTGAAGAACAAGGAA<br>GTGTCTGTAAAACGGGTTACCCAGGACCCTAAGCTCCAGATGGGCAAGAAGCTCCCGCTC<br>CACCTCACCCTGCCCCAGGCCTTGCCTCAGTATGCTGGCTCTGGAAACCTCACCCTGGCC<br>CTTGAAGCGAAAACAGGAAAGTTGCATCAGGAAGTGAACCTGGTGGTGATGAGAGCCACT<br>CAGCTCCAGAAAAATTTGACCTGTGAGGTGTGGGGACCCACCTCCCCTAAGCTGATGCTG<br>AGCTTGAAACTGGAGAACAAGGAGGCAAAGGTCTCGAAGCGGGAGAAGGCGGTGTGGGTG<br>CTGAACCCTGAGGCGGGGATGTGGCAGTGTCTGCTGAGTGACTCGGGACAGGTCCTGCTG<br>GAATCCAACATCAAGGTTCTGCCCACATGGTCGACCCCGGTGCAGCCAATGGCCCTGATT<br>GTGCTGGGGGGCGTCGCCGGCCTCCTGCTTTTCATTGGGCTAGGCATCTTCTTCTGTGTC<br>AGGTGCCGGCACTGA |
| **mClover3** | GTGAGCAAGGGCGAGGAGCTGTTCACCGGGGTGGTGCCCATCCTGGTCGAGCTGGACGGC<br>GACGTAAACGGCCACAAGTTCAGCGTCCGCGGCGAGGGCGAGGGCGATGCCACCAACGGC<br>AAGCTGACCCTGAAGTTCATCTGCACCACCGGCAAGCTGCCCGTGCCCTGGCCCACCCTC<br>GTGACCACCTTCGGCTACGGCGTGGCCTGCTTCAGCCGCTACCCCGACCACATGAAGCAG<br>CACGACTTCTTCAAGTCCGCCATGCCCGAAGGCTACGTCCAGGAGCGCACCATCTCTTTC<br>AAGGACGACGGTACCTACAAGACCCGCGCCGAGGTGAAGTTCGAGGGCGACACCCTGGTG<br>AACCGCATCGAGCTGAAGGGCATCGACTTCAAGGAGGACGGCAACATCCTGGGGCACAAG<br>CTGGAGTACAACTTCAACAGCCACTACGTCTATATCACGGCCGACAAGCAGAAGAACTGC<br>ATCAAGGCTAACTTCAAGATCCGCCACAACGTTGAGGACGGCAGCGTGCAGCTCGCCGAC<br>CACTACCAGCAGAACACCCCCATCGGCGACGGCCCCGTGCTGCTGCCCGACAACCACTAC<br>CTGAGCCATCAGTCCAAGCTGAGCAAAGACCCCAACGAGAAGCGCGATCACATGGTCCTG<br>CTGGAGTTCGTGACCGCCGCCGGGATTACACATGGCATGGACGAGCTGTACAAG |
| **mScarlet-I** | GTGAGCAAGGGCGAGGCAGTGATCAAGGAGTTCATGCGGTTCAAGGTGCACATGGAGGGC<br>TCCATGAACGGCCACGAGTTCGAGATCGAGGGCGAGGGCGAGGGCCGCCCCTACGAGGGC<br>ACCCAGACCGCCAAGCTGAAGGTGACCAAGGGTGGCCCCCTGCCCTTCTCCTGGGACATC<br>CTGTCCCCTCAGTTCATGTACGGCTCCAGGGCCTTCATCAAGCACCCCGCCGACATCCCC<br>GACTACTATAAGCAGTCCTTCCCCGAGGGCTTCAAGTGGGAGCGCGTGATGAACTTCGAG<br>GACGGCGGCGCCGTGACCGTGACCCAGGACACCTCCCTGGAGGACGGCACCCTGATCTAC<br>AAGGTGAAGCTCCGCGGCACCAACTTCCCTCCTGACGGCCCCGTAATGCAGAAGAAGACA<br>ATGGGCTGGGAAGCGTCCACCGAGCGGTTGTACCCCGAGGACGGCGTGCTGAAGGGCGAC<br>ATTAAGATGGCCCTGCGCCTGAAGGACGGCGGCCGCTACCTGGCGGACTTCAAGACCACC<br>TACAAGGCCAAGAAGCCCGTGCAGATGCCCGGCGCCTACAACGTCGACCGCAAGTTGGAC<br>ATCACCTCCCACAACGAGGACTACACCGTGGTGGAACAGTACGAACGCTCCGAGGGCCGC<br>CACTCCACCGGCGGCATGGACGAGCTGTACAAG |

**CFP-P2A-BlaR**

ATGGTGAGCAAGGGCGAGGAGCTGTTCACCGGGGTGGTGCCCATCCTGGTCGAGCTGGAC
GGCGACGTAAACGGCCACAAGTTCAGCGTGTCCGGCGAGGGCGAGGGCGATGCCACCTAC
GGCAAGCTGACCCTGAAGTTCATCTGCACCACCGGCAAGCTGCCCGTGCCCTGGCCCACC
CTCGTGACCACCCTGACCTGGGGCGTGCAGTGCTTCGCCCGCTACCCCGACCACATGAAG
CAGCACGACTTCTTCAAGTCCGCCATGCCCGAAGGCTACGTCCAGGAGCGCACCATCTTC
TTCAAGGACGACGGCAACTACAAGACCCGCGCCGAGGTGAAGTTCGAGGGCGACACCCTG
GTGAACCGCATCGAGCTGAAGGGCATCGACTTCAAGGAGGACGGCAACATCCTGGGGCAC
AAGCTGGAGTACAACGCCATCAGCGACAACGTCTATATCACCGCCGACAAGCAGAAGAAC
GGCATCAAGGCCAACTTCAAGATCCGCCACAACATCGAGGACGGCAGCGTGCAGCTCGCC
GACCACTACCAGCAGAACACCCCCATCGGCGACGGCCCCGTGCTGCTGCCCGACAACCAC
TACCTGAGCACCCAGTCCAAGCTGAGCAAAGACCCCAACGAGAAGCGCGATCACATGGTC
CTGCTGGAGTTCGTGACCGCCGCCGGGATCACTCTCGGCATGGACGAGCTGTACAAGGAA
TTCGGAAGCGGAGCTACTAACTTCAGCCTGCTGAAGCAGGCTGGAGACGTGGAGGAGAAC
CCTGGACCTCACGTGGCCAAGCCTTTGTCTCAAGAAGAATCCACCCTCATTGAAAGAGCA
ACGGCTACAATCAACAGCATCCCCATCTCTGAAGACTACAGCGTCGCCAGCGCAGCTCTC
TCTAGCGACGGCCGCATCTTCACTGGTGTCAATGTATATCATTTTACTGGGGGACCTTGT
GCAGAACTCGTGGTGCTGGGCACTGCTGCTGCTGCGGCAGCTGGCAACCTGACTTGTATC
GTCGCGATCGGAAATGAGAACAGGGGCATCTTGAGCCCCTGCGGACGGTGCCGACAGGTG
CTTCTCGATCTGCATCCTGGGATCAAAGCCATAGTGAAGGACAGTGATGGACAGCCGACG
GCAGTTGGGATTCGTGAATTGCTGCCCTCTGGTTATGTGTGGGAGGGCTAA

**LoxP Site**

ATAACTTCGTATAGCATACATTATACGAAGTTAT

**FrtF Site**

GAAGTTCCTATTCCGAAGTTCCTATTCTCTAGAAAGTATAGGAACTTC

**Frt3 Site**

GAAGTTCCTATTCCGAAGTTCCTATTCTTCAAATAGTATAGGAACTTC

**dClover2**

GTGAGCAAGGGCGAGGAGCTGTTCACCGGGGTGGTGCCCATCCTGGTCGAGCTGGACGGC
GACGTAAACGGCCACAAGTTCAGCGTCCGCGGCGAGGGCGAGGGCGATGCCACCATCGGC
AAGCTGACCCTGAAGTTCATCTGCACCACCGGCAAGCTGCCCGTGCCCTGGCCCACCCTC
GTGACCACCTTCGGCTACGGCGTGGCCTGCTTCAGCCGCTACCCCGACCACATGAAGCAG
CACGACTTCTTCAAGTCCGCCATGCCCGAAGGCTACGTCCAGGAGCGCACCATCTACTTC
AAGGACGACGGTACCTACAAGACCCGCGCCGAGGTGAAGTTCGAGGGCGACACCCTGGTG
AACCGCATCGAGCTGAAGGGCATCGACTTCAAGGAGGACGGCAACATCCTGGGGCACAAG
CTGGAGTACAACTTCAACAGCCACTACGTCTATATCACGGCCGACAAGCAGAACAACAGC
ATCAAGGCTAACTTCACCATCCGCCACAACGTTGAGGACGGCAGCGTGCAGCTCGCCGAC
CACTACCAGCAGAACACCCCCATCGGCGACGGCCCCGTGCTGCTGCCCGACAACCACTAC
CTGAGCCATCAGTCCGCCCTGAGCAAAGACCCCAACGAGAAGCGCGATCACATGGTCCTG
CTGGAGTTCGTGACCGCCGCCGGGATTACACATGGCATGGACGAGCTGTACAAG

**Supplementary Table S2: Sequences of single guide RNAs**

| Target | Guide sequence | Target sequence fw strand (PAM underlined) |
|---|---|---|
| CRY1 (fw strand) | GGAAACGTCCTAGTCAGGAAG | GGAAACGTCCTAGTCAGGAAGAGG |
| PER2-1 (rv strand) | CACCACCTGGTGTACCTCGC | CCAGCGAGGTACACCAGGTGGTG |
| PER2-2 (fw strand) | ATGGATCCCCCTTGAATCAC | ATGGATCCCCCTTGAATCACAGG |
| PER3-3 (fw strand) | GGCAGCCAGCGAGGTACACC | GGCAGCCAGCGAGGTACACCAGG |

**Supplementary Table S3: shRNA constructs**

| Target | Hairpin sequence (targeting sequence underlined) |
|---|---|
| PER2 (pGIPZ V2LHS_52938) | TGCTG TTGAC AGTGA GCGCG CATCC ATATT TCACT GTAAA<br>TAGTG AAGCC ACAGA TGTAT TTACA GTGAA ATATG GATGC<br>ATGCC TACTG CCTCG GA |
| CRY1 (pGIPZ V2LHS_172866) | TGCTG TTGAC AGTGA GCGCG CTGAG GCAAG CCGTT TGAAT<br>TAGTG AAGCC ACAGA TGTAA TTCAA ACGGC TTGCC TCAGC<br>ATGCC TACTG CCTCG GA |

**Supplementary Table S4: Sequences of PCR primers**

| Target | Sequence | Usage | Figure |
|---|---|---|---|
| CRY1 genomic locus (fw) | ACTGCCACTGATTGCCTGGGATTGAAGT | fw primer genomic PCR | Fig. S1D, Fig. S5C |
| CRY1 genomic locus (rv) | CAGCTGCAACAGTATTCCTCCTG | Rv primer Genomic PCR | Fig. S1D, Fig. S5C |
| PER2 genomic locus (fw) | ACCGGCTTCCAGGAGCCTCACTTGCA | fw primer genomic PCR | Fig. S1D, Fig. S5C |
| PER2 genomic locus (rv) | AAGCTGTCAGACTGAGTGGC | Rv primer genomic PCR | Fig. S1D, Fig. S5C |
| HF tag (rv) | TTGCTAGCCTTGTCGTCATC | RT primer | Fig. 1C Fig. S1C Fig. S5B |
| HF tag (rv) | ATCGTCTTTGTAGTCACCGCTACC | Rv primer RT-PCR | Fig. 1C Fig. S1C |
| CRY1 mRNA | TGCTGAGGCAAGCCGTTTGA | Fw primer RT-PCR | Fig. 1C Fig. S1C |
| PER2 mRNA | ACGCCCTTTCCACGTCAAGC | Fw primer RT-PCR | Fig. 1C Fig. S1C |
| mScarlet-I | GTCTTGAAGTCCGCCAGGTAGC | Rv primer RT-PCR | Fig. S5B |
| mClover3 | ACGCTGAACTTGTGGCCGTTT | Rv primer RT-PCR | Fig. S5B |

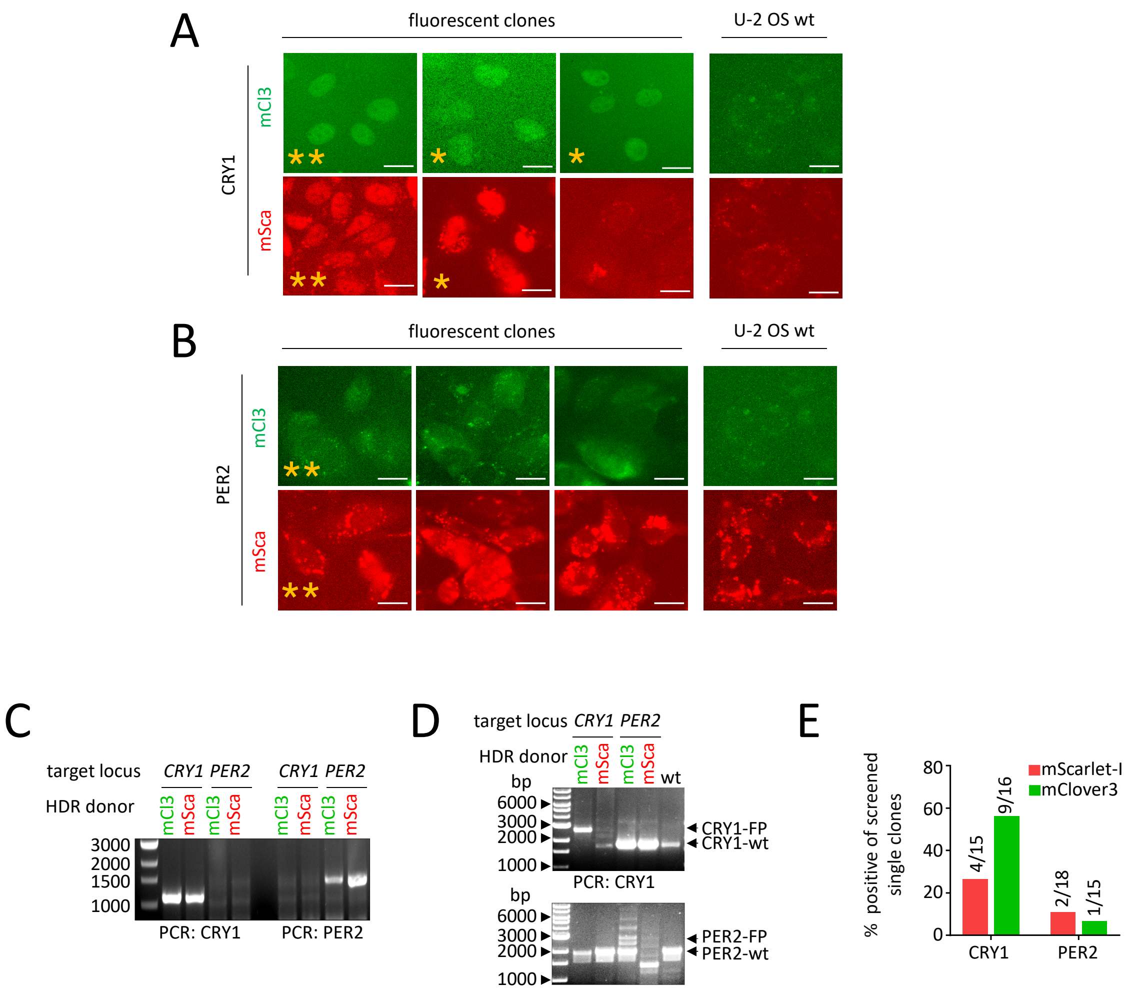


**Supplementary Figure S1: Screening of potential single knock-in clones. (A)** and **(B)**: Screening of potential CRY1 (A) and PER2 (B) knock-in clones by fluorescence microscopy using YFP and RFP channel. For each knock-in, 3 examples of fluorescent cells are shown along with wild-type cells that show autofluorescence only. Clones that were confirmed positive for correct knock-in by PCR afterwards are marked with *, clones used for all further analysis with **. **(C)** Chimeric mRNA was detected in single clones by RT-PCR (as in Fig. 1C). **(D)** Successful knock-in was confirmed by amplification of the targeted region of PER2 and CRY1 genomic loci by PCR, which shows either wild-type allele, the larger knock-in allele or both. Additional bands are probably heteroduplexes of wild-type and knock-in product. **(E)** Percentage of positive knock-in clones in relation to all screened clones for the 4 different knock-in experiments. Numbers indicate count of correct and initially screened clonal colonies, respectively.

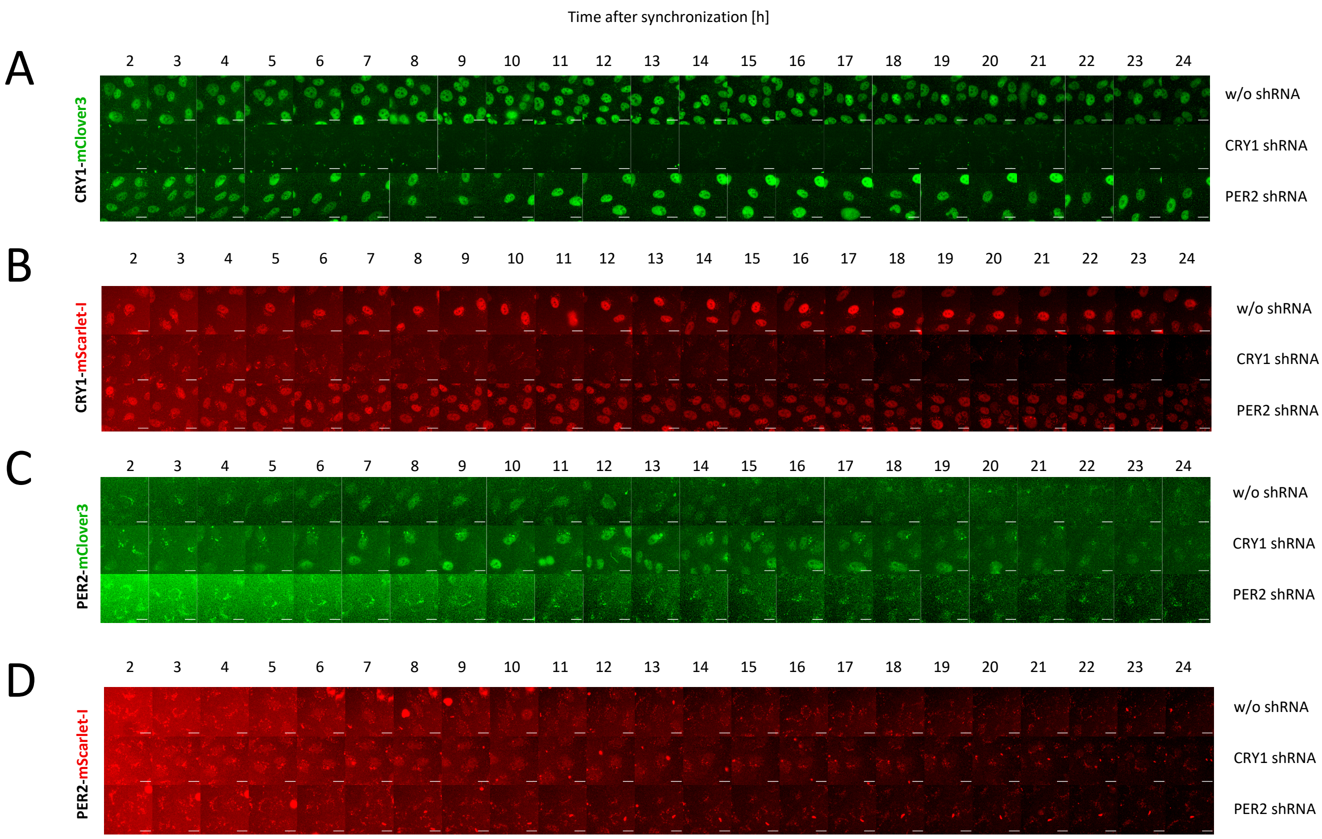


**Supplementary Figure S2**: **Complete time series of knock-down experiment (Fig. 1F).** U-2 OS knock-in cells expressing CRY1-mClover3 **(A)**, CRY1-mScarlet-I **(B)**, PER2-mClover3 **(C)** or PER2-mScarlet-I **(D)**, respectively, were either left untreated or transduced with shRNA targeting either CRY1 or PER2. After synchronization, fluorescence in the respective channel was recorded for 24 hours. Scale bar: 20 µm.

Supplementary Figure S2

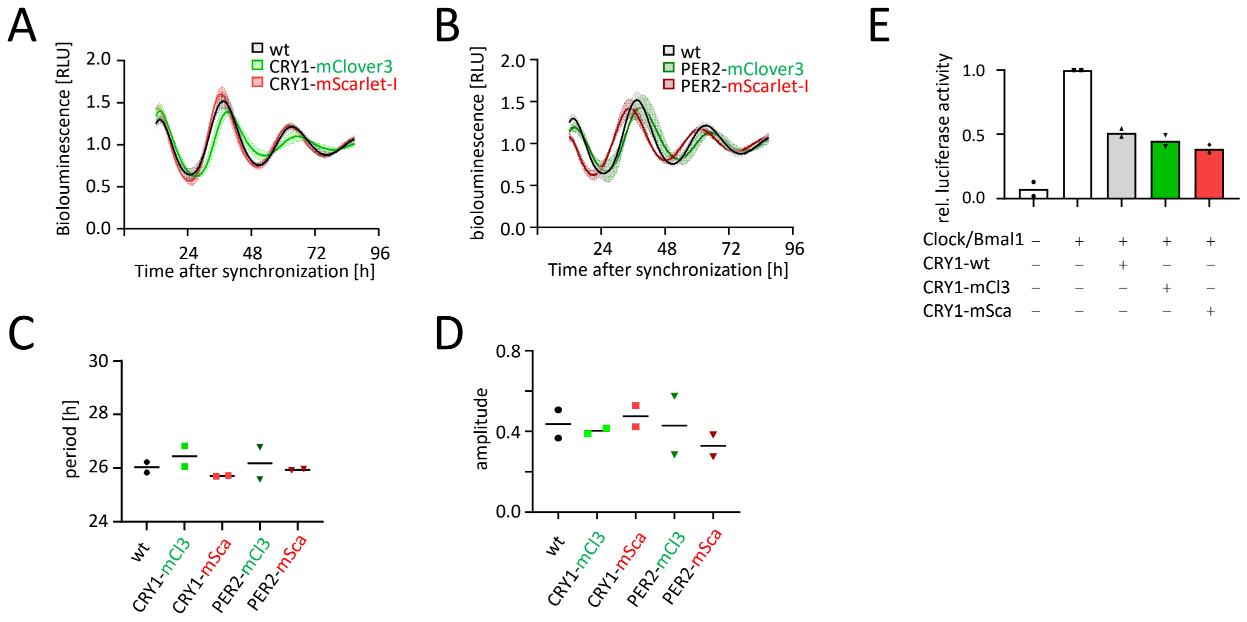


**Supplementary Figure S3: Analysis of circadian rhythms in single knock-in cells. (A)-(D)** Individual clones and wild-type cells were transduced with a *Bmal1*:Luc reporter and luminescence was recorded over four days. Depicted are mean + SD of four individual, detrended traces resulting from two independent experiments **(A)** and **(B)**, and mean calculated period lengths **(C)** and amplitude **(D)** for both experiments. **(E)** Ability of CRY fusion proteins to inhibit CLOCK/BMAL1 induced activation of an E-BOX reporter plasmid. HEK-293 cells were transfected with an 6xEBOX-Luciferase reporter plus the indicated constructs and reporter activity was measured (n=2).

Supplementary Figure S3

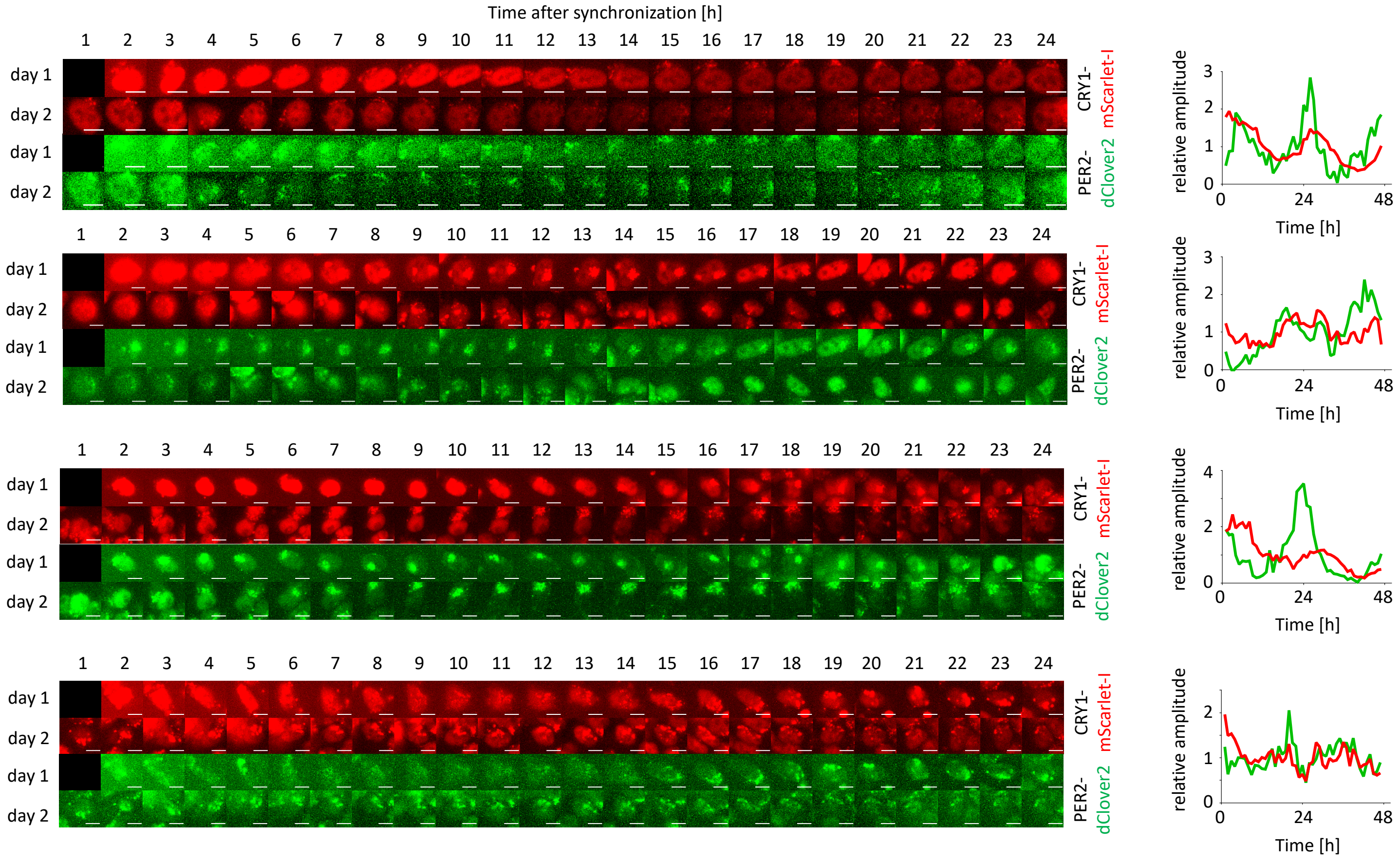


**Supplementary Figure S4: Time series of HCT-116 double knock-in cells.** Montages of bicolor fluorescence microscopy images of individual HCT-116 double-knock-in (PER2-dClover2/CRY1-mScarlet-I) cell's nuclei over the course of 2 days. Traces of 4 individual cells are shown. Mean nuclear fluorescence signals were quantified, backgrounds subtracted and signals normalized by dividing by mean signal of the time course. Scale bar: 10 µM.

Supplementary Figure S4

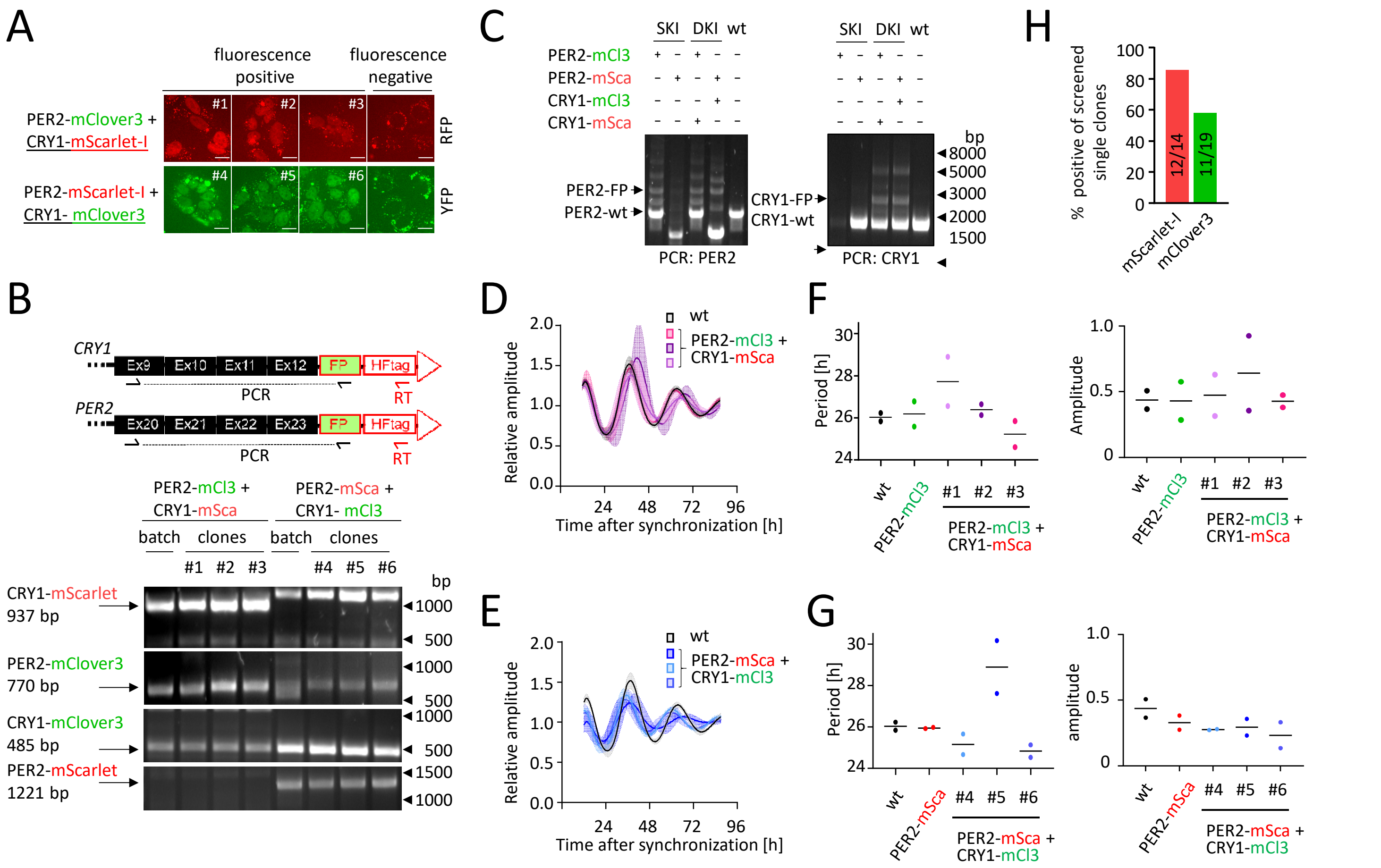


**Supplementary Figure S5: Selection and characterization of double knock in clones. (A)** Screening of clones with potential CRY1-knock in by fluorescence microscopy. For each knock-in, 3 example clones with the expected pattern are shown along with a negative clone. **(B)** Chimeric mRNA was detected in the three single clones from A by RT-PCR using RT-primer specific to the insertion, gene specific forward and fluorophore specific reverse primer. Arrows indicate the expected band for righteous insertion. **(C)** Successful knock-in was confirmed by amplification of the edited genomic locus using out-out PCR followed by Sanger sequencing. Results exemplarily shown for DKI clones #3 and #6. **(D-G)** Individual double knock-in clones, the corresponding parental clone and wt cells were transduced with a *Bmal1*:Luc reporter and luminescence was recorded over four days. Depicted are mean + SD of four individual, detrended traces resulting from two independent experiments **(D-E)**, and mean calculated period lengths and amplitude for both experiments **(F-G)**. Clone #6 was used for imaging analysis. **(H)** Percentage of positive knock-in clones in relation to all screened clones. Scale bar: 20 µm. mCl3 = mClover3, mSca = mScarlet-I, FP = fluorescent protein (mScarlet-I or mClover3).

### Supplementary Figure S5
